# Deciphering transcriptomic signatures explaining the phenotypic plasticity of non-heading lettuce genotypes under artificial light conditions

**DOI:** 10.1101/2023.02.01.526118

**Authors:** Hiroto Yamashita, Kaede C. Wada, Noritoshi Inagaki, Zui Fujimoto, Jun-ichi Yonemaru, Hironori Itoh

## Abstract

Elucidating the mechanisms and pathways involved in genotype–environment (G×E) interactions and phenotypic plasticity is critical for improving plant growth. Controlled environment agricultural systems allow growers to modulate the environment for particular genotypes. In this study, we evaluated the effects of interactions among 14 genotypes and four artificial light environments on leaf lettuce phenotypes and dissected the underlying molecular mechanism via transcriptome-based modeling. Variations in morphological traits and phytochemical contents in response to artificial light treatments revealed significant G×E interactions. The appropriate genotype and artificial light combinations for maximizing phenotypic expression were determined on the basis of a joint regression analysis and the additive main effect and multiplicative interaction model for these G×E interactions. Transcriptome-based regression modeling explained approximately 50%–90% of the G×E variations. Further analyzes indicated *Red Lettuce Leaves 4 (RLL4*) regulates UV-B and blue light signaling through the effects of the HY5–MBW pathway on flavonoid biosynthesis and contributes to natural variations in the light-responsive plasticity of lettuce traits. Our study represents an important step toward elucidating the phenotypic variations due to G×E interactions in non-heading lettuce under artificial light conditions.

**Highlights:**

- Several morphological characteristics of lettuce genotypes were altered by different light wavelengths.
- A defective *RLL4* allele (*rll4*) induces the expression of downstream genes related to UV-B and blue light signaling through activation of the HY5–MBW pathway, which enhances phytochemical accumulation in lettuce.
- G×E analyzes identified the ideal genotype and artificial light combinations for individual phenotypes.
- Transcriptome-based modeling explained approximately 50%–90% of the G×E variations.

## Introduction

Crop production in the future will need to be more sustainable to minimize the effects of the environment (Fedoroff *et al*. 2010; West *et al*. 2014). There are increasing risks and uncertainties associated with crop production in fields because of climate change and related extreme weather events. Controlled environment agricultural systems, such as closed-type plant factories and vertical farms, enable growers to control several environmental factors, including light sources, temperature, CO_2_ concentration, and nutrient composition and content (Shamshiri *et al*. 2018). Thus, enclosed systems facilitate risk-free and year-round crop production because there are no variable factors affecting production or seasonal effects that are associated with field production. Although controlled environment agricultural production is becoming increasingly feasible and may be important for ensuring the cultivation of nutrient-rich food crops (Pinstrup-Andersen 2018), it typically relies on legacy cultivars that have been selected and bred for field conditions (Folta 2019). Optimizing cultivars is essential for increasing crop productivity and nutritional performance under enclosed artificial conditions.

Lettuce (*Lactuca sativa* L.), which is one of the most important vegetable crops consumed worldwide, has nutritional benefits because it is a source of dietary fiber, several important dietary minerals, various vitamins, and bioactive phytochemicals (e.g. carotenoids and phenolics) (Kim *et al*. 2016). Common lettuce phenolics are caffeic acid derivatives (predominantly chicoric, chlorogenic, caffeoyltartaric, and caffeoylmalic acids) and flavonol glycosides (predominantly quercetin-glycosides) (Llorach *et al*. 2008; Oh *et al*. 2009). Red-type lettuce cultivars contain cyanidin-glycosides (anthocyanins) (Llorach *et al*. 2008).

Lettuce cultivars vary substantially in terms of morphological characteristics and phytochemical contents (Llorach *et al*. 2008; Kim *et al*. 2016; Zhang *et al*. 2017, 2022; Yang *et al*. 2018). In a previous study, RNA-seq and bulked segregant analyzes were performed to clone four *Red Lettuce Leaves* genes (*RLL1* to *RLL4*), which are responsible for the variations in the red coloration of lettuce (Su *et al*. 2020). The *RLL1* and *RLL2* genes encode a bHLH (basic helix-loop-helix) transcription factor and an R2R3-MYB transcription factor, respectively. The MYB, bHLH, and WD40-repeat transcription factors combine to form a complex (MBW) that mainly regulates the expression of flavonoid biosynthetic pathway genes (Gonzalez *et al*. 2008; Xu *et al*. 2015). In a large-scale RNA-seq analysis of 240 *Lactuca* spp. genotypes, a genome-wide association study (GWAS) identified 12 expression quantitative trait loci (eQTLs) that regulate flavonoid biosynthesis-associated genes and several QTLs controlling anthocyanin accumulation (Zhang *et al*. 2017). The accumulation of lettuce phytochemicals is sensitive to environmental conditions, including temperature and light (intensity and quality) (Oh *et al*. 2009; Becker *et al*. 2014; Kitazaki *et al*. 2018; Assumpção *et al*. 2019). Recent studies used narrow-band light emitting diodes (LEDs) to determine the effects of different artificial light qualities and intensities on the leaf lettuce transcriptome and/or metabolome, with a particular focus on phytochemicals (Kitazaki *et al*. 2018; Nagano *et al*. 2022). These omics-based analyzes clarified the coordinated regulation of flavonoid accumulation and the expression patterns of biosynthesis-associated genes in lettuce exposed to various LED sources. Lettuce morphology also changes in response to various light conditions (Kim *et al*. 2004). However, most of these previous studies examined a single environment or genotype (e.g., only comparing genotypes in one environmental condition or evaluating the effects of various light sources on a single genotype). There have been relatively few studies that evaluated genotype–environment (G×E) interactions during responses to light.

Light is an essential energy source for plant photosynthesis and is also a key signal for initiating morphogenesis and metabolic changes. There is a wide variety of photoreceptors in higher plants. For example, phytochromes perceive red/far-red light (600–750 nm), whereas cryptochromes, phototropins, and F-box-containing flavin-binding proteins perceive blue/UV-A light (320–500 nm) and UV RESISTANCE LOCUS 8 (UVR8) perceives UV-B light (280– 320 nm) (Paik & Huq 2019). ELONGATED HYPOCOTYL 5 (HY5) is a key transcription factor that regulates the plant light response, which is activated by phytochromes, cryptochromes, and UVR8 via the dissociation of HY5 from the CONSTITUTIVELY PHOTOMORPHOGENIC 1 (COP1)/SUPPRESSOR OF PHYA-105 (SPA) complex (Gangappa & Botto 2016). Among the HY5-induced genes, *REPRESSOR OF UV-B PHOTOMORPHOGENESIS 1 (RUP1*) and *RUP2* encode WD40-repeat proteins that function as negative feedback regulators of UVR8 (Gruber *et al*. 2010). The BLUE-LIGHT INHIBITOR OF CRYPTOCHROMES (BIC), which bind directly to cryptochromes and inhibits their active dimerization (Wang *et al*. 2016), and RUPs function as negative feedback circuit regulators of blue light and UV-B signals (Gruber *et al*. 2010; Wang *et al*. 2017; Tissot & Ulm 2020). The *RLL4*, which contributes to the variable red coloration of lettuce, encodes a *RUP* gene that encodes a negative regulator of UV-B signaling (Su *et al*. 2020). However, the effects of light quality (e.g., UV-B light) on *RLL4* alleles remain unknown.

Our group reported that different combinations of *RLL* alleles result in quantitative differences in the accumulation of phytochemicals, especially anthocyanins, among lettuce cultivars under artificial light conditions (Wada *et al*. 2022). Although the genetic variation has been confirmed, the differences due to multiple artificial light environments have not been demonstrated. The present study was designed to clarify the G×E interactions affecting the non-heading lettuce response to artificial light. Specifically, we evaluated the G×E interactions for 14 genotypes and four artificial light conditions. The statistical analysis of the G×E interactions identified the genotype and artificial light environment combinations that optimized leaf morphological traits and phytochemical contents. Furthermore, transcriptome-based modeling dissected the molecular and metabolic dynamics.

## Materials and Methods

### Plant materials and artificial light treatments

In this study, 14 non-heading lettuce (var. *crispa* or var. *longifolia*) genotypes were used, of which nine and five were red-type and green-type cultivars, respectively (Table S1). Red-Batavia (RButt), Red-Leaf (RLeaf), Red-Oak (ROak), Red-Coslettuce (RCos), Fancy-Red (FancR), Handsome-Red1 (HandR), Banchu-Redfire (RFire), Sun-Bright (SunB), and Wine-Dress (WineD) were the red-type cultivars, whereas Green-Oak (GOak), Handsome-Green (HandG), Green-Leaf (GLeaf), Green-Batavia (GBat), and Frill-Ice (FrIce) were the green-type cultivars. Seeds were obtained from Nakahara Seed Product Co., Ltd. (Fukuoka, Japan), Yokohama Nursery Co., Ltd. (Kanagawa, Japan), and Takii & Co., Ltd. (Kyoto, Japan).

Seeds were soaked with distilled water on a urethane sponge in a plastic cell tray (4 cm × 4 cm/cell) and pre-germinated at 22 °C under a white LED (16-h light/8-h dark cycle; 180 μmol m^-2^ s ^-1^ PPFD). After 1 week, the plants were transferred to a half-strength Otsuka House A-recipe nutrient solution (pH 5.2) (OAT Agrio Co., Ltd., Tokyo, Japan) and then incubated under the same light conditions as before. After 10 days, the plants were weighed and exposed to the following four artificial light conditions: white LED (HLU-E0_N23; Dai Nippon Printing Co., Ltd., Tokyo, Japan), red LED (660 nm), blue LED (445 nm), and white fluorescent light (FHF32EX-N-HX-S; NEC Corporation, Tokyo, Japan). The red LED and blue LED treatments were performed using a three-in-one LED (3LH-75DPS; Nippon Medical and Chemical Instruments, Osaka, Japan). The fluorescent light included approximately 20 μW/cm^2^ UV light (260–395 nm), as determined using a UV intensity meter (CENTER532 ST; Sato Shoji Inc., Kanagawa, Japan). In contrast, UV intensity was undetectable for the white LED, red LED, and blue LED. The spectra for the light conditions are presented in Figure S1. After a 3-day light treatment, the shoots were harvested, weighed, and immediately frozen in liquid nitrogen and stored at −80 °C prior to the HPLC analysis and RNA extraction. Growth during the 3-day treatment period was assessed on the basis of the increase in the fresh weight.

The following eight cultivars were used for the UV-B assay: RLeaf, ROak, FancR, HandR, WineD, GLeaf, GButt, and FrIce. Plants were pre-cultured as described above and then exposed to the white LED (180 μmol m^-2^ s^-1^ PPFD) (HLU-E0_N23; Dai Nippon Printing Co., Ltd, Tokyo, Japan) with or without UV-B (approximately 20–30 μW/cm^2^) provided by the UV-LED irradiation module (UV-SPOT 310 nm; THORLABS, Inc., Tokyo, Japan). Plants were irradiated with UV for 16 h during the day and the daytime temperature was maintained at 22 °C.

The time-course study of the response to white fluorescent light was completed using the following four cultivars: RLeaf, FancR, GLeaf, and FancG. Plants were pre-cultured as described above and then incubated under white fluorescent light (120 μmol m^-2^ s^-1^ PPFD) (FHF32EX-N-HX-S; NEC Corporation, Tokyo, Japan). Samples were harvested at 0, 6, 24, and 48 h after initiating the exposure to white fluorescent light.

### Plant phenotyping according to non-destructive 3D imaging

A non-destructive 3D imaging analysis was performed using an all-surroundings imaging system (Kochi *et al*. 2018; Kochi *et al*. 2021; Kochi *et al*. 2022). The system consisted of four cameras (TRI089S-CC, resolution 4096 × 2160, CMOS sensor; LUCID Vision Labs Inc., Richmond B.C., Canada) and LED lighting. Each plant was placed on a turntable and rotated by five degrees using a motor to capture images from all directions. Our original closed-loop coarse-to-fine method (Hayashi *et al*. 2020) was used to estimate the camera position, whereas a modified Structure-from-motion/multi-view stereo method was used to process the captured images. The measurement accuracy of this system was less than 1 mm (Kochi *et al*. 2022). The number of expanded leaves was determined using the constructed 3D images. The following parameters were measured for three representative expanded leaves and the average values were calculated: leaf length, leaf width, and the length to width ratio.

### Quantification of lettuce phytochemical contents

The relative contents of the following nine phytochemicals in lettuce extracts were determined: chicoric acid, chlorogenic acid, three cyanidin-glycosides [cyanidin 3-*O*-(3-*O*-malonyl-beta-D-glucoside) (Cy3_3MG), cyanidin 3-*O*-(6-*O*-malonyl-beta-D-glucoside) (Cy3_6MG), and cyanidin 3-*O*-glucoside (Cy3G)], and four quercetin-glycosides [quercetin-3-*O*-(6-*O*-malonyl)-beta-D-glucoside-7-*O*-glucuronide (Q3_6MbGG), quercetin-3-*O*-(6”-*O*-malonyl)-glucoside-7-*O*-glucoside (Q3_6MGG), quercetin-3-*O*-(6-*O*-malonyl)-glucoside (Q3_6MG), and quercetin-3-*O*-glucuronide/quercetin-3-*O*-glucoside (Q3G)]. Previously described methods that were modified slightly were used for the analysis (Arapitsas *et al*. 2008; Wada *et al*. 2022). Powdered lettuce samples were extracted with methanol-water formic acid solution (40:59:1, v/v/v) for 1 h after 3 min sonication, followed by two extractions with 80% (v/v) methanol. The 1100 Series HPLC system (Agilent Technologies, Waldbronn, Germany) equipped with a YMC-Triart C18 column (150 mm × 2.0 mm I.D., S-3 μm, 12 nm; YMC Co., Ltd., Kyoto, Japan) was used in this study. A 10 μl aliquot of the extract was injected into the HPLC system. The flavonoids and phenolic acids in the extract were separated using a mobile phase (0.2 ml min^-1^) consisting of a 5% (v/v) formic acid aqueous solution (A) and acetonitrile (B). The elution was performed with the following gradient at 28 °C: initial concentration of 5% B, followed by a 3 min hold at 5% B, 2 min linear gradient from 5% to 10% B, 3 min linear gradient from 10% to 12% B, 2 min linear gradient from 12% to 14% B, 8 min hold at 14% B, 10 min linear gradient from 14% to 17% B, 7 min linear gradient from 17% to 20% B, 5 min linear gradient from 20% to 90% B, 5 min hold at 90% B, 5 min linear gradient from 90% to 5% B, and 8 min hold at 5% B. The compounds in the eluent, excluding cyanidin-glycosides, were detected at 350 nm, while the cyanidin-glycosides were monitored at 520 nm. The phenolic compounds were identified according to the HPLC data (UV spectra and retention times) and an analysis using an electrospray ionization tandem mass spectrometry (ESI-MS/MS) system (UltiMate 3000 RSLC system; Thermo Fisher Scientific, Waltham, MA, United States). Compounds were quantified using calibration curves for the following chemicals: chicoric acid and chlorogenic acid purchased from Toronto Research Chemicals (Toronto, Canada), quercetin-3-*O*-glucoside purchased from Extrasynthese (Genay, France), and cyanidin 3-*O*-glucoside purchased from Nagara Science (Gifu, Japan). The abundances of cyanidin derivatives and quercetin derivatives were calculated as the equivalent amount of quercetin-3-*O*-glucoside and cyanidin 3-*O*-glucoside, respectively.

### G×E interaction analysis

The G×E interactions for phenotypes were evaluated by a two-way analysis of variance (ANOVA), a joint regression analysis (JRA) (Finlay & Wilkinson 1963), and an analysis involving the additive main effect and multiplicative interaction (AMMI) model (Gauch 1988).

For the JRA, the Finlay-Wilkinson model was as follows: *y_ij_* = *g_i_* + *b_i_ x_j_* + *e_ij_*, where *y_ij_* is the phenotypic value of genotype *i* in environment *j, g_i_* is the genetic effect (intercept), *b_i_* is the regression coefficient (slope indicates stability), *x_j_* is the environmental index value (average phenotypic value of all genotypes in environment *j*), and *e_ij_* is the residual.

The AMMI model was as follows: 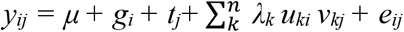, where *y_ij_* is the phenotypic value of genotype *i* in environment *j*, *g_i_* is the genotype *i* mean deviation (genotype mean minus grand mean), *t_j_* is the environment *j* mean deviation (environment mean minus grand mean), *λk* is the singular value for the interaction principal component (IPC) axis *k, u_ki_* is the genotype *i* eigenvector value for the IPC axis *k*, *v_kj_* is the environment *j* eigenvector value for the IPC axis *k*, and *e_ij_* is the residual. The biplot graph of the first and second IPC axes was visualized using the R package “ggplot2” (ver. 3.3.6).

### RNA-seq analysis

Total RNA was extracted using the ISOSPIN Plant RNA kit (Nippon Gene, Tokyo, Japan). The RNA concentration was measured using the Qubit fluorometer (Thermo Fisher Scientific, Waltham, MA, United States). The mRNA was purified using oligo-dT beads (New England Biolabs Japan Inc., Tokyo, Japan) and then the sequencing library was constructed using the NEBNext Ultra II Directional RNA Library Prep Kit for Illumina (New England Biolabs Japan Inc., Tokyo, Japan). The library was sequenced using the illumina NovaSeq 6000 platform with S4 flow cells to produce 150 bp paired-end and dual index reads. The RNA-seq data were deposited in the DDBJ Sequence Read Archive (accession number: DRA014848).

Reads were preprocessed using Trimmomatic (ver. 0.39) with the following parameters: ILLUMINACLIP TruSeq3-PE-2.fa:2:30:10, LEADING:15, TRAILING: 15, SLIDINGWINDOW:10:15, and MINLEN:50. After the preprocessing step, the remaining reads were mapped to the lettuce reference genome (Reyes-Chin-Wo *et al*. 2017) using STAR (ver. 2.7.8a) (Dobin *et al*. 2013). Gene expression levels were estimated using RSEM (ver. 1.3.3) (Li & Dewey 2011). The RSEM output was subsequently analyzed using the R software (ver. 4.0.2) (R Core Team 2020). After excluding genes with average counts <10, the count data were normalized against the trimmed mean of M values (TMM) and the library size using the R package “edgeR” (ver.3.30.3). The data were log2-transformed and used for the subsequent analyzes.

A principal component analysis (PCA) was performed using the R “prcomp” function and the first and second principal components (PC1 and PC2) were visualized. The “glmLRT” function in the R package “edgeR” (ver.3.30.3) was used for the likelihood ratio test. Genes with a false discovery rate (FDR) < 0.05 were extracted as the differentially expressed genes among all groups. These genes were included in the *k*-means cluster analysis, which was completed using the “*k*-means” function in the R package. The appropriate number of clusters (*k* = 6) was determined according to the elbow method (Figure S2). Heatmaps and UpSet plots were prepared using the R package “ComplexHeatmap” (ver. 2.11.1). For the Gene Ontology (GO) enrichment analysis, the eggNOG mapper was used to annotate the lettuce reference genome with GO terms (Cantalapiedra *et al*. 2021). The GO enrichment analysis was performed using the “runTest” function in the R package “TopGO” (ver. 2.40.0) and the following options: algorithm = “elim” and statistic = “fisher”. The genes used for the pathway expression analyzes are summarized in Table S2.

### Regression modeling

For the regression modeling, phenotypic values and the transcriptome data (TMM-normalized count data) were used as the response and explainable variables, respectively. A flow chart for the modeling is presented in Figure S3. All data were divided into a training (70%) and test (30%) dataset using a stratified sampling approach for the red-type and green-type lettuce cultivars. Noninformative transcriptome data were eliminated using the Boruta feature selection algorithm (Kursa & Rudnicki 2010). More specifically, the following parameters of the “Boruta” function in the R package Boruta (ver. 7.0.0) were applied: getImp = getImpRfZ, maxRuns = 1000, and pValue = 0.01. This feature selection process was repeated 100 times to ensure the results were robust. For the regression modeling, genes selected 10–100 or more times (at intervals of 10) in 100 repetitions were included in the informative transcriptome. Regression models were then generated using the informative transcriptome and Random Forest algorithms (method = “rf”) in the R package “caret” (ver. 6.0-92). The hyperparameter “mtry” was optimized via a random search involving a repeated cross-validation with the following parameters: number = 10, repeats = 3, and tuneLength = 10. To assess the accuracy of predictions, the coefficient of the determination (R^2^), root-mean-square error, and mean absolute error were calculated for the observed and predicted values. The black-box model was interpreted according to the variable importance and partial dependence plot (pdp). The variable importance was calculated using the “varImp” function in the R package “caret” (ver. 6.0-92). The pdp was produced using the “partial” function in the R package “pdp” (ver. 0.8.0).

## Results

### Phenotypic variations among lettuce genotypes under artificial light conditions

We evaluated the effects of the interactions among 14 genotypes and four artificial lights (white LED, red LED, blue LED, and fluorescent light) on lettuce phenotypic variations. Several morphological traits of lettuce genotypes were altered by the exposure to different light wavelengths (Figure 1a, b). With the exception of leaf number, the morphological traits revealed G×E interactions (Figure S4; two-way ANOVA, FDR < 0.05). For the red light treatment, many cultivars varied regarding leaf length and width (Figure 1b). The analysis of nine phytochemicals (chicoric acid, chlorogenic acid, three cyanidin-glycosides, and four quercetin-glycosides) in lettuce extracts also revealed G×E interactions (Figure S4; two-way ANOVA, FDR < 0.05). The cyanidin-glycoside and quercetin-glucoside contents increased significantly in nine red-type genotypes under blue light and fluorescent light conditions, but the content changes differed markedly among the cultivars (Figure 2a). The Q3_6MbGG and Q3G contents in five green-type genotypes also increased following the fluorescent light treatment (Figure 2a). The G×E interactions were weak for the chicoric acid and chlorogenic acid contents (Figure S4), which increased in most genotypes exposed to white light and blue light conditions, respectively (Figure 2a).

**Figure 1.**
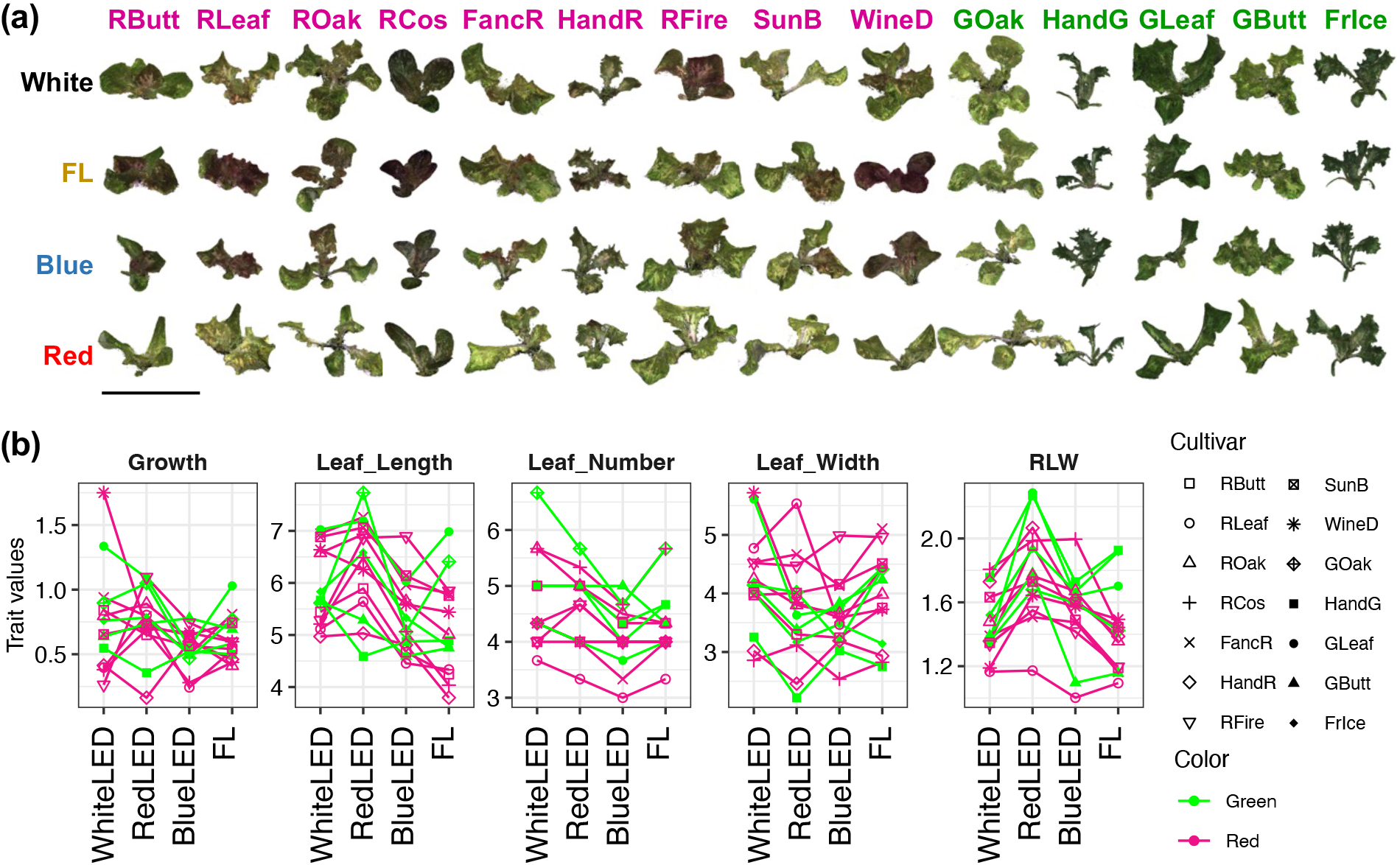
Morphological phenotypic variations of leaf lettuce cultivars under artificial light conditions. **(a)** 3D reconstructed images of leaf lettuce cultivars under four artificial light conditions. Bar = 10 cm. **(b)** Interaction plots of morphological phenotypic values. Plots with different shapes are mean in each cultivar. Light magenta and green lines are red-type (nine) and green-type (five) cultivars, respectively.

**Figure 2.**
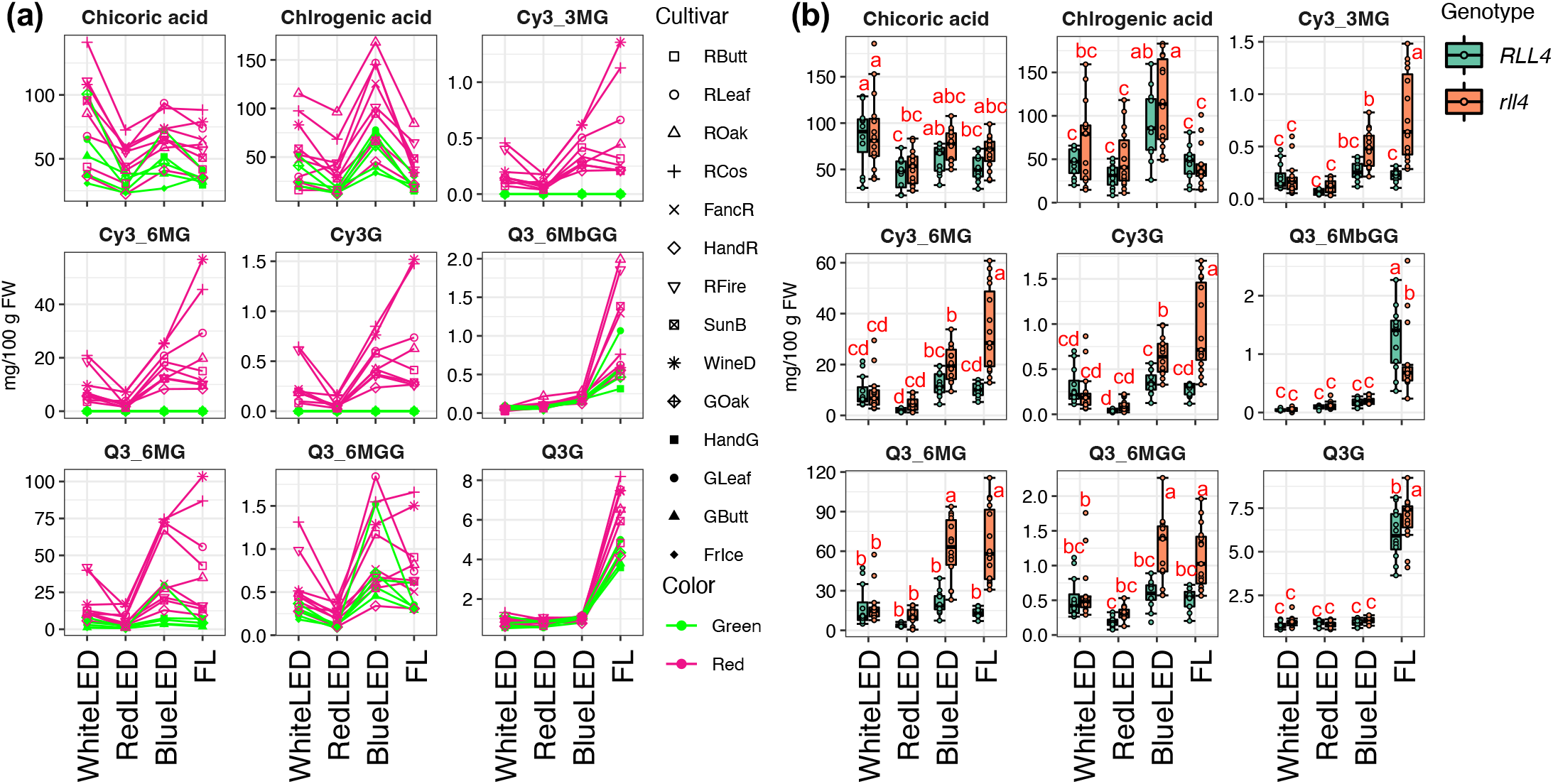
Phytochemical variations of leaf lettuce cultivars under artificial light conditions. (**a)** Interaction plots of phytochemical contents. Plots with different shapes are mean in each cultivar. Light magenta and green lines are red-type (nine) and green-type (five) cultivars, respectively. **(b)** Boxplots of phytochemical contents among *RLL4* alleles in red-type cultivars. Different letters indicate significant differences (Tukey’s HSD test, *P* < 0.05).

In the previous study, we genotyped *RLL* genes and anthocyanidin synthase (*ANS*)genes in 133 leaf lettuce accessions, including the 14 genotypes used in this study. Our analysis confirmed that three functional alleles for *RLL1*, *RLL2*, and *ANS* form the core gene set responsible for red coloration and a defective *RLL4 (rll4*), which is an ortholog of *RUP* that encodes a negative regulator of UV-B signaling in *Arabidopsis*, enhances anthocyanin accumulation under fluorescent light (Wada *et al*. 2022). In the current study, we compared the phytochemical contents of lettuce cultivars carrying different *RLL4* alleles under four artificial light conditions. The cyanidin-glycoside and quercetin-glycoside contents in the *rll4* genotypes increased substantially under blue and fluorescent lights (Figure 2b).

To evaluate the temporal changes in phytochemical metabolism, we cultivated two red-type (RLeaf and FancR) and two green-type (GLeaf and FancG) lettuce genotypes under fluorescent light and collected samples at 0, 6, 24, and 48 h. The FancR and RLeaf red-type genotypes carried the *RLL4* and *rll4* alleles, respectively. In the RLeaf samples, the nine examined phytochemicals accumulated quickly after 6 h and continued to increase until 48 h (Figure S5a). In the FancR samples, chicoric acid, chlorogenic acid, and quercetin-glycoside contents increased slightly after 6 h and cyanidin-glycosides accumulated at 48 h (Figure S5a). In the green-type cultivars GLeaf and FancG with *RLL4*, the phytochemicals accumulated slightly after 6 h, with the exception of the cyanidin-glycosides (Figure S5a).

The fluorescent light used in this study included the UV wavelength. Thus, we also examined the responses of eight lettuce genotypes to UV-B using a white LED and a UV irradiation module. The UV-B treatment considerably increased most of the phytochemical contents in the *rll4* genotypes (Figure 3a, b), which was relatively consistent with the observed changes under fluorescent light conditions.

**Figure 3.**
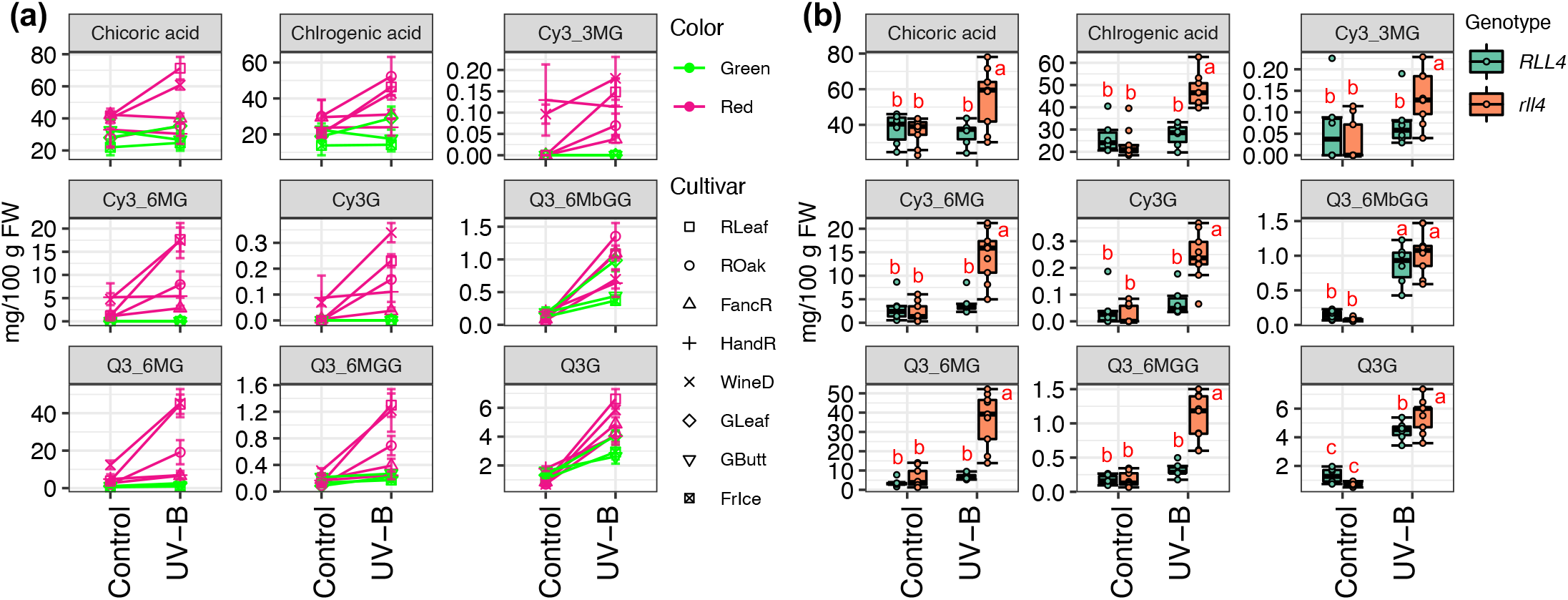
Effects of UV-B radiation in phytochemical contents of leaf lettuce eight cultivars. (**a)** Interaction plots of phytochemical contents. Plots with different shapes and error bars are mean±SD in each cultivar. Light magenta and green lines are red-type (five) and green-type (three) cultivars, respectively. **(b)** Boxplots of phytochemical contents among *RLL4* alleles in red-type cultivars. Different letters indicate significant differences (Tukey’s HSD test, *P* < 0.05)

### Analysis of G×E interactions among lettuce phenotypes

To determine which combinations of genotypes and artificial light environments were ideal for lettuce cultivation, we analyzed the G×E interactions for morphological traits and phytochemicals using the JRA and AMMI models. The JRA quantified the phenotypic plasticity of each genotype under artificial light conditions (Figures 4a, b, S6, 7). For the WineD, RLeaf, RCos, ROak, and RButt cultivars (defective *RLL4* allele), the regression coefficient (*b*: stability) of the Finlay-Wilkinson model was high (*b* > 1) for the cyanidin-glycoside and quercetin-glycoside contents (Cy3_6MG and Q3_6MG) (Figure 4b). The stability values were also high for the chlorogenic acid contents of RLeaf, RCos, ROak, FancR, and HandG and the chicoric acid contents of GOak, RCos, RFire, SunB, WineD, ROak, and GLeaf (Figure 4a, b). These results reflect the quantitative differences in the G×E variations among 14 lettuce genotypes and indicate appropriate combinations of genotypes and artificial light conditions may enhance phytochemical accumulation in lettuce. In addition, the AMMI model was used for a qualitative analysis of the phenotypic plasticity of each genotype under artificial light conditions. The constructed biplot indicated the genotype and artificial light environment combinations that produced desirable phenotypes (Figures 4c, S6, 7).

**Figure 4.**
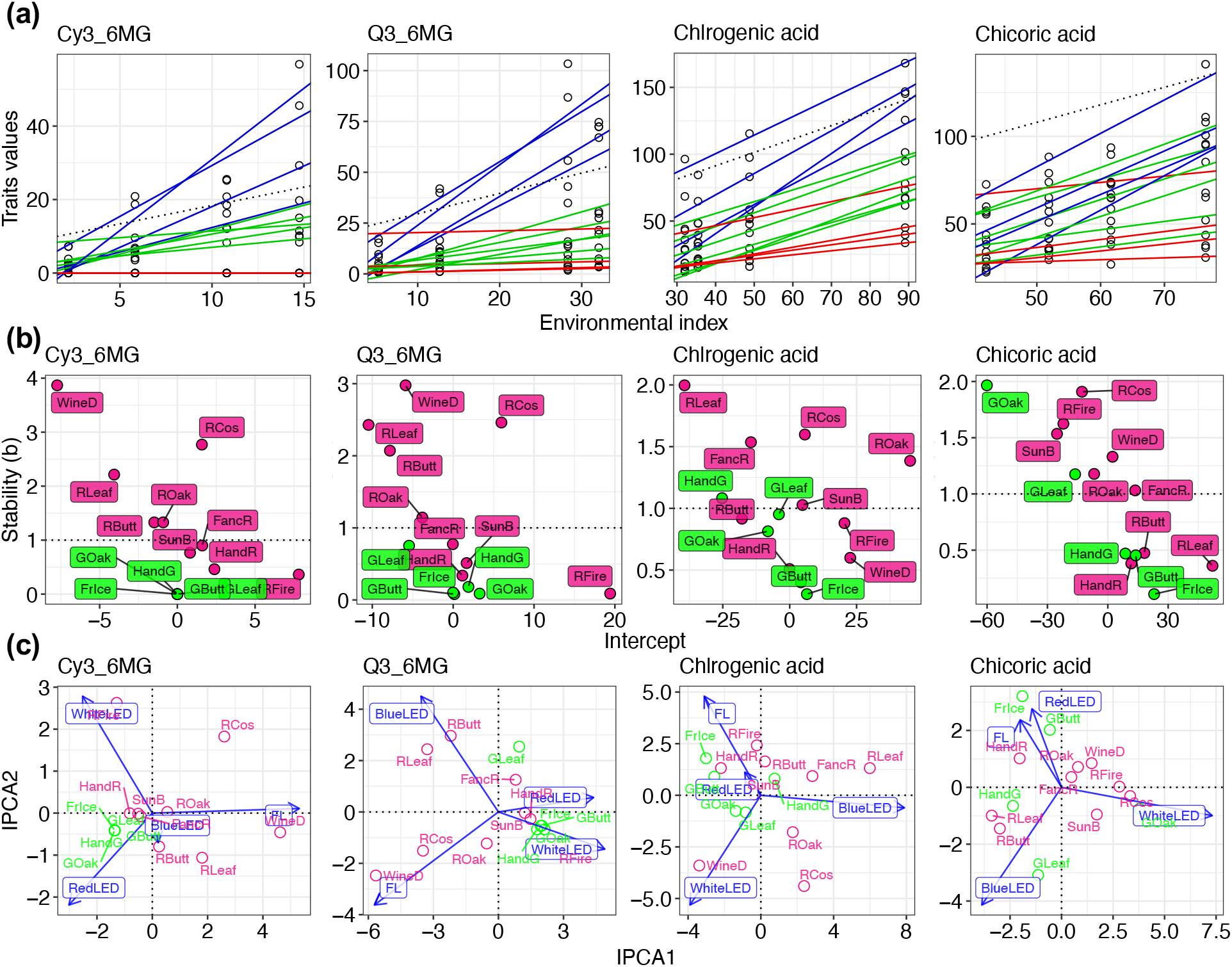
The analysis of G×E interaction for leaf lettuce phenotypes. **(a, b)** The joint regression analysis (JRA) based on Finlay-Wilkinson model. **(a)** Each plot indicates the observed phenotypic value for each genotype in an artificial light environment, and each line is the regression-fitted value for each genotype. **(b)** The relationship of each stability (b), means each slope, and intercept in the regression-line for each genotype. **(c)** Biplots of additive main effects and multiplicative interaction (AMMI) model. The first and second interaction principal component axes (IPCA) were plotted. Analyses for representative four metabolites were shown in this figure. The results of the other metabolites were also shown in Figure S6.

### Transcriptomic dissections of G×E variations

To dissect the molecular mechanism underlying the phenotypic plasticity of each genotype in response to artificial light environments, we analyzed the transcriptomes of 165 shoot samples from 14 genotypes exposed to four artificial light conditions. We obtained a mean of 11 million filtered fragments (read pairs) per sample (Table S3). The PCA indicated the transcriptomes were diverse, with their profiles roughly separated in terms of lettuce color types and artificial light conditions (Figure 5a). The transcriptomes of the red-type and green-type genotypes were classified into six clusters (RC01–RC06 and GC01–GC06; Figure 5b, c). Many of these light source-specific expressed genes were common to the red-type and green-type genotypes (Figure 5d). The RC02 and GC03 clusters contained the blue light-specific expressed genes. The enriched GO terms among these genes included “chloroplast organization” (GO: 0009658), “thylakoid membrane organization” (GO:0010027), and “Group II intron splicing” (GO:0000373) (Figure 5b–e, Table S4). In contrast, the enriched GO terms assigned to the red light-specific expressed genes in the RC04 and GC02 clusters, included “microtubule-based movement” (GO:0007018), “cortical microtubule organization” (GO:0043622), and “plant-type secondary cell wall biogenesis” (GO:0009834) (Figure 5b–e, Table S4).

**Figure 5.**
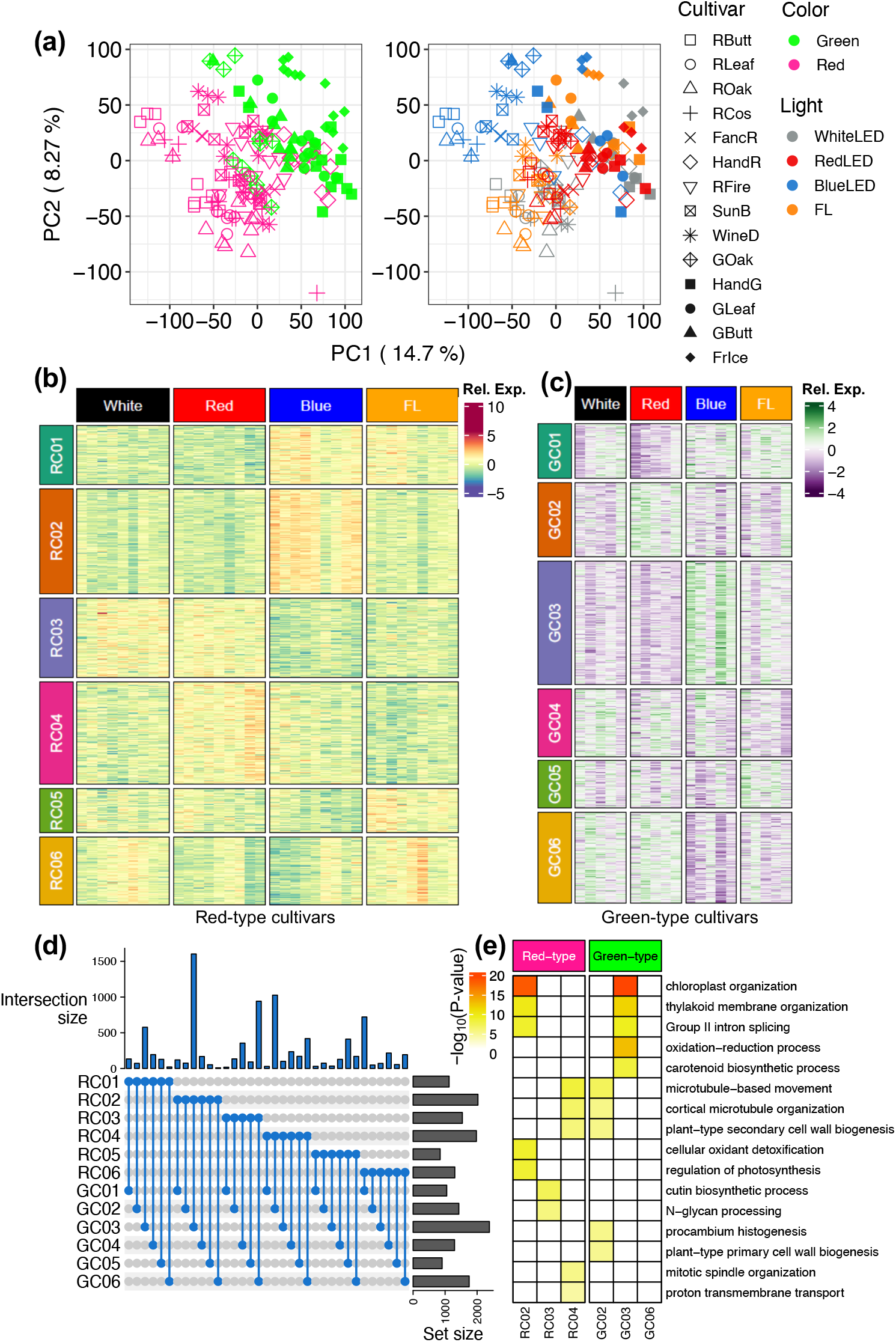
Global transcriptomic variation of leaf lettuce cultivars under artificial light conditions. **(a)** Principal component analysis (PCA) of transcriptomes. The left and right figures show the different coloring based the lettuce color types and artificial light treatments, respectively. PC1 and PC2 were plotted. **(b, c)** Heatmaps of transcriptome profiles in red-type and green-type lettuce cultivars. **(d)** UpSet plot of genes in each cluster. **(e)** GO enrichment analysis for genes in clusters 2, 3, and 4 in red-type and cluster 2, 3, and 6 in green-type of **(b, c)**. –log_10_ p-values of five top-ranked GO term in each cluster were shown as heatmaps.

To estimate the extent to which the transcriptome can explain the phenotypic variations due to G×E interactions, we constructed transcriptome-based regression models for lettuce phenotypes. First, the noninformative transcriptome data were removed using the Boruta feature selection algorithm. For the phytochemical analysis, 2–20 genes were selected in all 100 repetitions (Figure 6a, Table S5). The expression patterns of these genes explained approximately 60%–90% of the phenotypic variations (Figure 6b). The predictions for the cyanidin-glycoside contents were the most accurate, with the expression patterns of 18–19 genes explaining almost all of the phenotypic variation (Figure 6a, b). The expression of only five and two genes explained approximately 60% of the phenotypic variations in the chicoric acid and chlorogenic acid contents, respectively (Figure 6a, b). In terms of the morphological traits, the prediction accuracy decreased after 70 or more genes were selected (Figure 6a, b). Accordingly, we extracted 70 or more selected genes as key informative genes, of which 6–19 genes explained approximately 40%–60% of the phenotypic variations (Figure 6a, b, Table S6). Some of the Boruta-selected genes for the phytochemical contents were common among metabolites and included enzyme-encoding genes (e.g., *CHS, CHI, F3H, F3’H, FLS, DFR*, and *ANS*) and R2R3-MYB transcription factor genes involved in flavonoid metabolism (Figure 6c). The Boruta-selected genes for morphological traits were generally not shared among the traits and included auxin signaling pathway genes [e.g., *AUXIN RESPONSE FACTOR (ARF*)and *Auxin/INDOLE-3-ACETIC-ACID (Aux/IAA)]* affecting the leaf width and length (Figure 6c).

**Figure 6.**
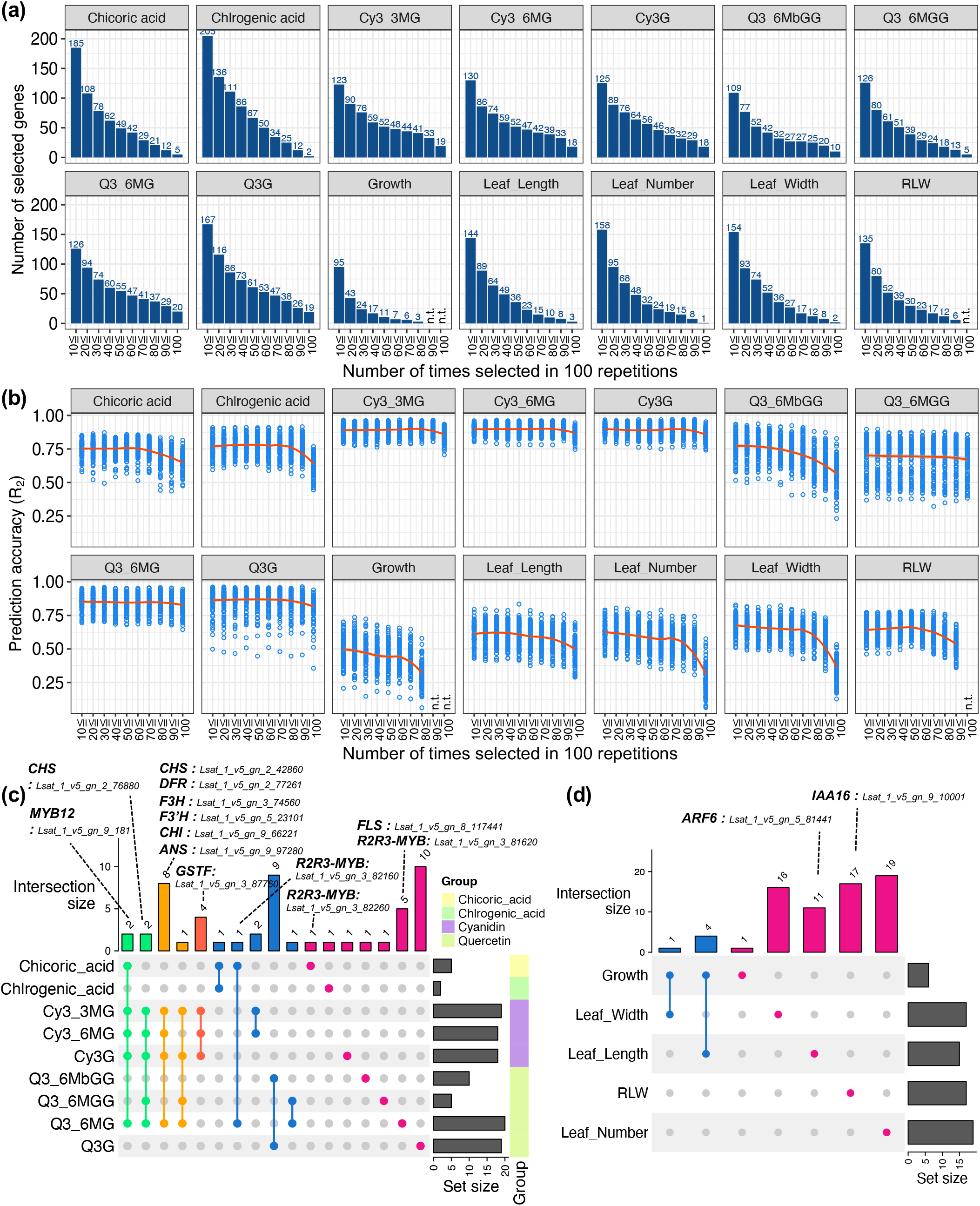
Feature selection and the performance of transcriptome-based modeling. **(a)** The number of genes selected by a Boruta feature selection. Genes of 10-100 or more times (at 10 intervals) out of 100 repetitions in each phenotype were shown. n.t.: not tested. **(b)** Prediction accuracy of the transcriptome-based modeling based on Boruta-selected genes. Plots show the results in each repetition, and the red curves show LOESS smoothing lines. **(c)** UpSet plot of Boruta-selected genes of all repetitions in lettuce phytochemicals. **(d)** UpSet plot of Boruta-selected genes of 70 times out of 100 in morphological phenotypes.

The variable importance and pdp in the regression models constructed using the random forest algorithm enabled the interpretation of nonlinear relationships between phenotypes and gene expression levels (Figures 7a–d, S8). The pdp of the phytochemical contents gradually increased as the gene expression levels increased (Figures 7c, d, S8). We confirmed the G×E interactions for the expression of genes affecting phenylpropanoid and flavonoid metabolism. Many of these genes had light wavelength-responsive expression patterns that were linked to phenotypic plasticity (Figures 2a, 8a). In addition, the blue and fluorescent light treatments substantially increased the expression of these genes in the genotypes with a defective *RLL4* allele (Figure 8b). Furthermore, the examination of the temporal response to fluorescent light conditions and the UV-B treatment indicated the expression patterns of these genes were related to phenotypic plasticity (Figures S5b, 9).

**Figure 7.**
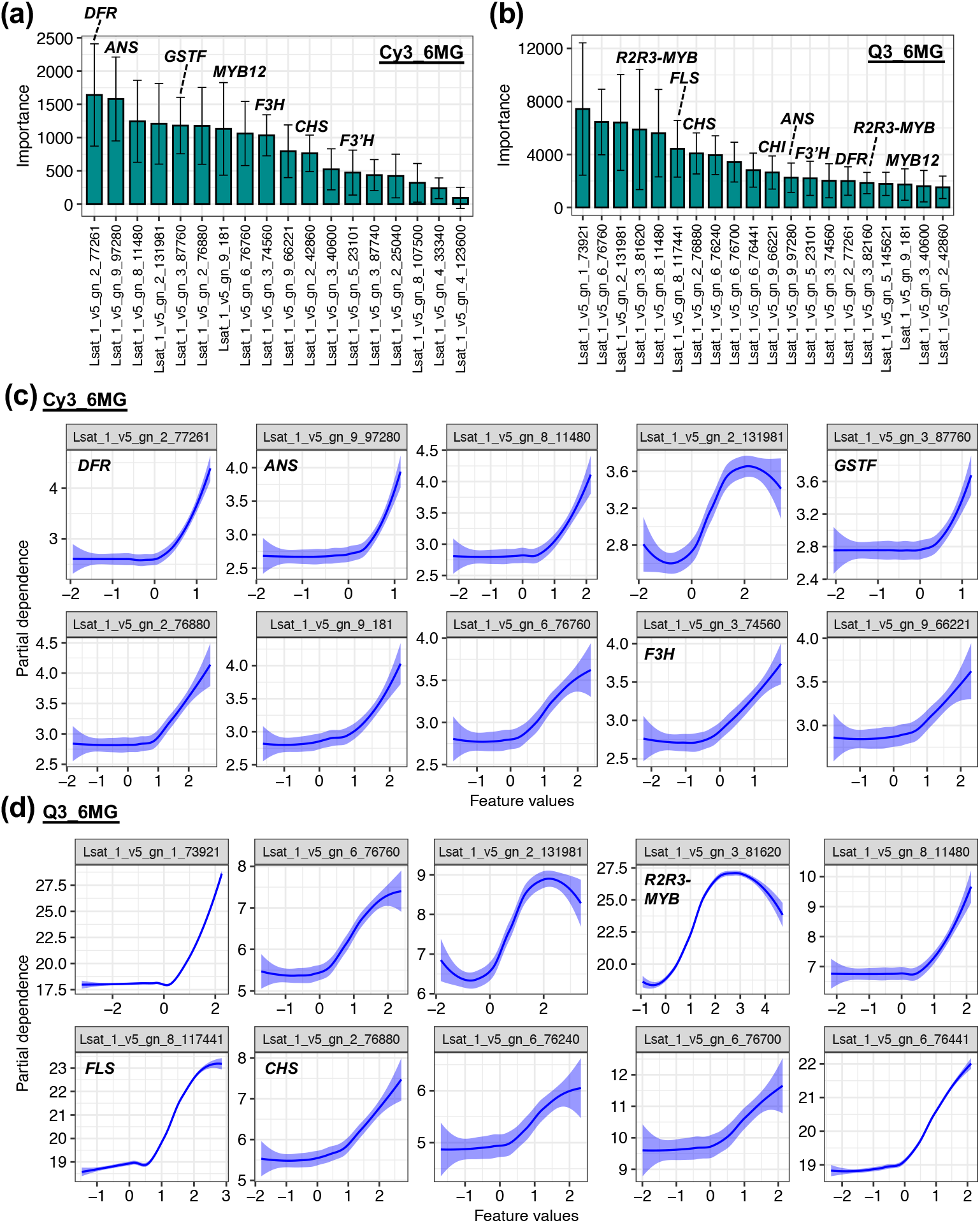
Interpreting of a transcriptome-based black-box model. **(a, b)** Variable importance in the regression model based on random forest algorithm for cyanidin 3-*O*-(6-*O*-malonyl-beta-D-glucoside) (Cy3_6MG) and quercetin-3-*O*-(6-*O*-malonyl)-glucoside (Q3_6MG) contents. Data and error bars are the mean±SD (100 repetitions). **(c, d)** Partial dependence plots (pdp) in the regression model based on random forest algorithm for Cy3_6MG and Q3_6MG contents. Blue curves show LOESS smoothing lines of the pdp in 100 repetitions.

**Figure 8.**
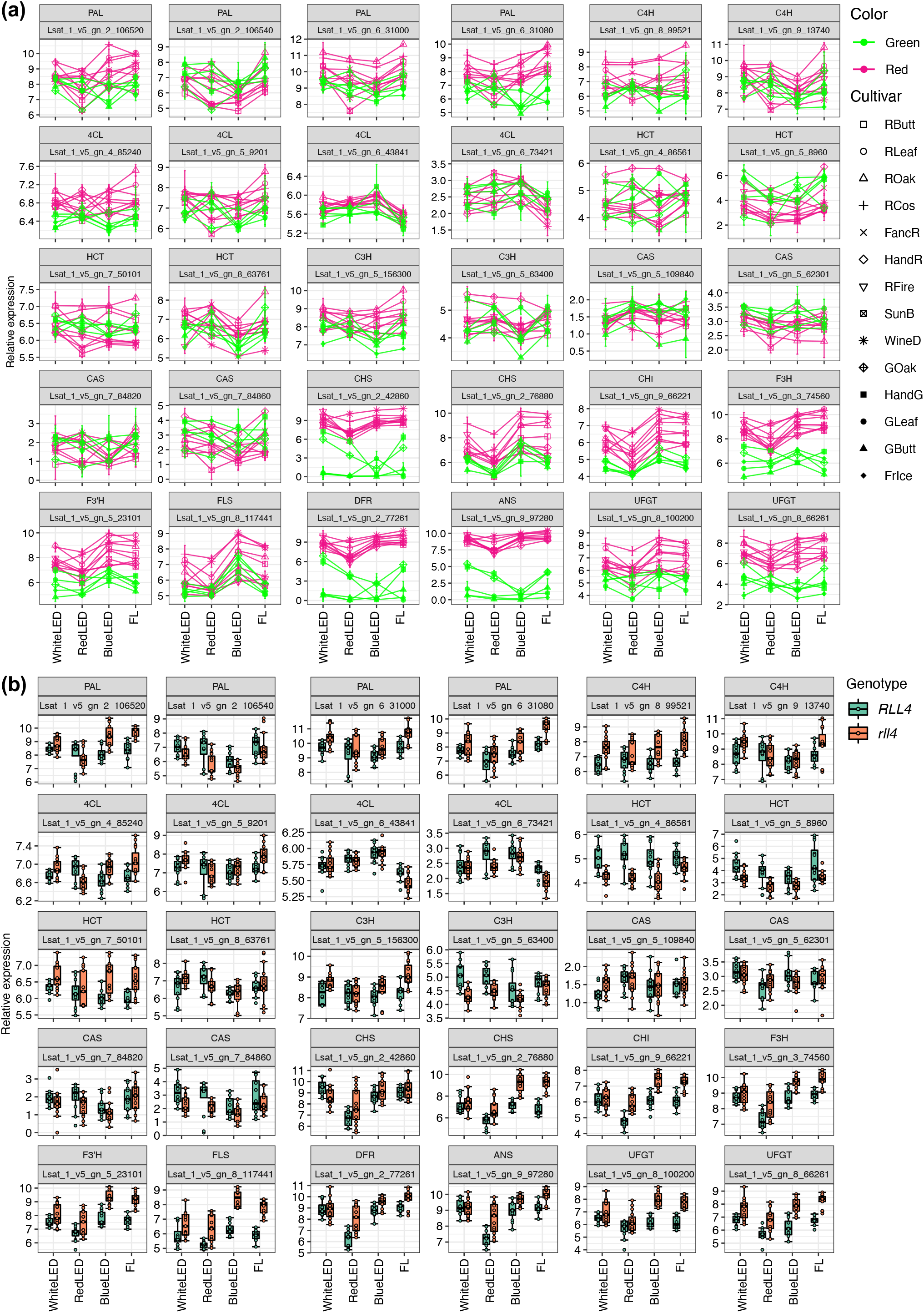
Expression patterns of phenylpropanoid and flavonoid pathway genes under artificial light conditions. **(a)** Interaction plots of expression levels. **(b)** Boxplots of expression levels among *RLL4* alleles in red-type cultivars.

## Discussion

The importance of controlled environment agricultural systems for crop production will likely increase because they are unaffected by global climate change (Pinstrup-Andersen 2018). However, field-bred cultivars are typically used in these systems. To identify the optimal cultivars for enclosed artificial settings, the mechanisms and physiological pathways associated with G×E interactions and phenotypic plasticity must be elucidated. In this study, we assessed the effects of the interactions involving 14 genotypes and four artificial lights (white, red, and blue LEDs and fluorescent light) on lettuce phenotypes by analyzing morphological traits, phytochemical contents, and transcriptome data.

Diverse morphological characteristics and phytochemical contents in 14 lettuce genotypes were responsive to light wavelengths (Figures 1, 2). The cyanidin-glycoside and quercetin-glucoside contents in nine red-type genotypes increased significantly, but differentially among the lettuce genotypes, under blue light, fluorescent light, and UV-B-supplemented conditions (Figures 2a, 3a). In most plants, including lettuce, flavonoid compounds accumulate in response to UV-B and blue light because they can absorb the corresponding wavelength and they function as reactive oxygen species scavengers (García-Macías *et al*. 2007; Kitazaki *et al*. 2018; Ferreyra, Serra & Casati 2021). Thus, the results of the current study suggest that lettuce has developed a light-induced flavonoid accumulation mechanism that facilitates adaptations to various light-stress environments. A previous study identified four genes (*RLL1*–*RLL4*) that modulate the anthocyanin (cyanidin-glycoside) content in lettuce (Su *et al*. 2020). Moreover, we previously showed that a mutated *RLL4* allele (*rll4*), which is a *RUP* gene related to the negative feedback regulation of UV-B signaling (Gruber *et al*. 2010; Su *et al*. 2020), promotes cyanidin-glycoside and quercetin-glucoside accumulation under fluorescent lights (Wada *et al*. 2022). Therefore, we compared these phytochemical contents among lettuce cultivars with different *RLL4* alleles under four artificial light and UV-B-supplemented conditions. The cyanidin-glycoside and quercetin-glycoside contents in the *rll4* genotypes increased dramatically under blue light, fluorescent light, and UV-B-supplemented conditions (Figures 2b, 3b). These findings suggest that RLL4 may repress blue light and UV-B signaling in lettuce, although RUP1 acts as a repressor for UV-B response in *Arabidopsis* (Gruber *et al*. 2010).

Phenylpropanoid and flavonoid metabolism-associated gene expression in the genotypes with the *rll4* allele also increased considerably under blue light, fluorescent light, and UV-B-supplemented conditions (Figures 7b, S5b, S9b). In plants, HY5 is a major transcription factor regulating light signaling and the expression of downstream genes associated with light acclimation and tolerance, including genes related to flavonoid biosynthesis and photoprotection (Gangappa & Botto 2016). The *HY5* expression levels in the genotypes with the *rll4* allele increased in response to the blue light, fluorescent light, and UV-B-supplemented conditions (Figure S10). These light conditions also increased the expression levels of MBW complex genes, especially the R2R3 MYB (*Lsat_1_v5_gn_3_82160, Lsat_1_v5_gn_3_82260*, and *Lsat_1_v5_gn_3_81620*) and bHLH (*Lsat_1_v5_gn_5_190001*)transcription factor genes, in the cultivars carrying the *rll4* allele (Figure S11). In *Arabidopsis*,HY5 regulates *PAP1* (R2R3-MYB) expression by binding directly to the promoter; *PAP1* expression is highly dependent on HY5 (Shin *et al*. 2013). The *RUP* and *BIC* expression levels are up-regulated downstream of HY5 via the cross-regulation of blue light and UV-B signaling, respectively, and the encoded proteins function as negative feedback circuit regulators (Gruber *et al*. 2010; Wang *et al*. 2017; Tissot & Ulm 2020). Artificial light conditions also increased the expression of the lettuce *RLL4* and *BIC* genes and other genes (e.g., *COP1, SPA1*, and *CRY2*) in the genotypes with *rll4* (Figure S12). These results indicate that a defect in *RLL4* leads to the induced expression of downstream genes associated with UV-B and blue light signaling via the increased expression of the *HY5* and MBW-related genes, thereby enhancing phytochemical accumulation in lettuce. These findings imply that the *RLL4* allele generally contributes to the natural variations in the responses of lettuce accessions to light.

The feature selection and regression modeling in this study were completed using the phenotypes and transcriptomes of 165 shoot samples in four artificial light environments. Transcriptome-based regression modeling explained approximately 60%–90% and 40%–60% of the G×E variations in the phytochemical contents and morphological traits, respectively (Figure 6). Hence, the changes in the transcriptome profiles and metabolite contents are closely linked. However, the model performance was lower for the morphological traits than for the phytochemical contents (Figure 6). One explanation for this difference is that the plasticity of the changes in the leaf length and width was not fully captured because the treatment period was short (3 days). The transcriptomes in this study were for entire shoot samples. Accordingly, the relationship between morphological changes and the transcriptome may be more precisely determined if only the leaves are used for phenotyping or analyzes of temporal alterations.

Our transcriptome modeling may have captured the molecular differences in the light-induced changes in phytochemical metabolites. Compared with the other quercetin-glycosides and cyanidin-glycosides, the light treatments differentially affected Q3G and Q3_6MbGG (Figures 2a, 3a). When the *RLL4* allele was defective (*rll4*), almost all of the tested flavonoids accumulated in response to UV-B and blue light. However, the UV-B-responsive accumulation of Q3G and Q3_6MbGG occurred independently of the *RLL4* allele (Figures 2b, 3b). In addition, these two metabolites were unaffected by blue light regardless of the *RLL4* allele (Figure 2b). During the regulated accumulation of Q3G and Q3_6MbGG, there may be metabolic regulatory mechanisms in addition to the network associated with the expression of *RLL4* and the downstream genes. This possibility is supported by the unique key informative genes selected during the transcriptome modeling for Q3G and Q3_6MbGG (Figure 6c, Table S5). The selected genes included several superoxide dismutase genes (*Lsat_1_v5_gn_6_16060, Lsat_1_v5_gn_6_43340*, and *Lsat_1_v5_gn_8_10740*), suggestive of a possible relationship between flavonoid accumulation and antioxidant activity. Nevertheless, the diversity in the accumulation of flavonoid-glycosides in response to blue light or UV-B wavelengths remains to be comprehensively investigated.

Almost all of the lettuce phenotypes evaluated in this study revealed G×E interactions (Figure S4). To determine which combination of genotype and artificial light environment is most conducive to lettuce production, we analyzed our data using JRA and AMMI models. In breeding programs, the AMMI model is useful for identifying the best performing cultivars in specific environments (Gauch 1988). These G×E analyzes have been conducted for several field crops (Huang *et al*. 2016; Li *et al*. 2021), but they have rarely been performed for genotypes in various artificial environments. Furthermore, these G×E statistical approaches may be relevant for optimizing cultivar performance in controlled environment agricultural systems. Specifically, the environmental conditions may be customized according to the results of these analyzes. In the present study, the ideal genotypes and artificial light environments for each phenotype were determined. For example, the anthocyanin (Cy3_6MG) content was highest in WineD incubated under fluorescent light (Figure 3c). These results will help breeders select lettuce cultivars with desirable morphological and phytochemical traits in controlled environment agricultural systems.

The present study dissected the G×E interactions for lettuce phytochemicals and morphological traits under artificial light conditions on the basis of statistical analyzes and transcriptome modeling. The AMMI analysis revealed the optimal combinations of genotypes and artificial light environments for lettuce traits. Notably, growing lettuce genotypes with a defective *RLL4* allele under blue light or UV-B-supplemented conditions may substantially increase lettuce phytochemical quality. Our transcriptome data were used to clarify the variations in these G×E interactions. For example, flavonoid accumulation was mainly influenced by the diversity in the related gene expression levels. These light-induced changes to gene expression may serve as useful biomarkers when selecting for G×E effects prior to phenotypic examinations. Our study represents an important step toward elucidating the genotype-dependent phenotypic plasticity of several crops under artificial light conditions. The information provided herein will be applicable for selecting and breeding cultivars suitable for controlled environment agricultural systems.

## Acknowledgments

We thank N. Kochi and A. Hayashi for their assistance with the non-destructive 3D imaging analysis. We thank Y. Ogo and T. Kawakatsu for helping to construct the cDNA library for the RNA-seq analysis. We thank Y. Nakai for helpful discussions regarding this project. We thank M. Noda and K. Nakatani for their technical assistance during experiments. Lettuce seeds were kindly provided by Nakahara Seed Product Co., Ltd. (Japan) and Yokohama Nursery Co., Ltd. (Japan). We thank the Research Center for Advanced Analysis of NARO for the use of facilities. We thank Edanz (https://jp.edanz.com/ac) for editing a draft of this manuscript.

