## Supplementary_Figures for "Deciphering transcriptomic signatures explaining the phenotypic plasticity of non-heading lettuce genotypes under artificial light conditions"

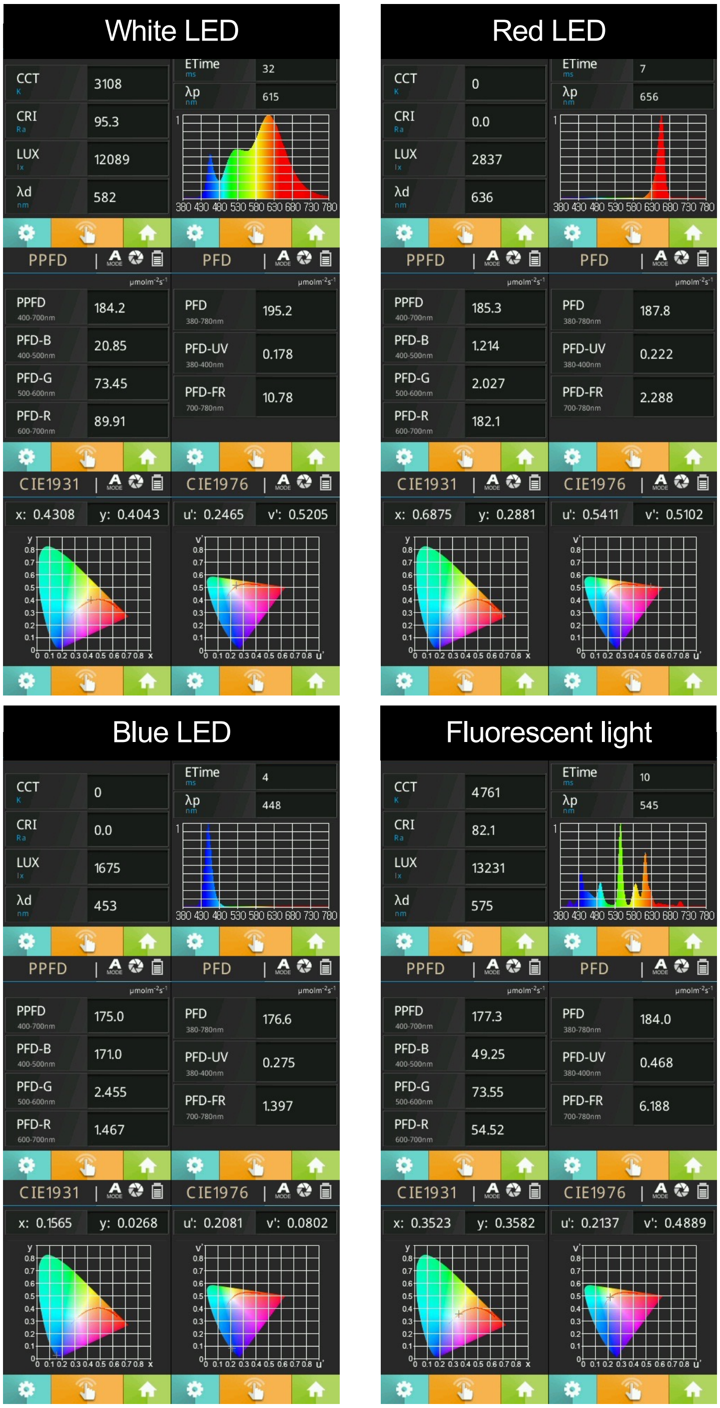


**Figure S1 Spectra information of four artificial light conditions in this study.**

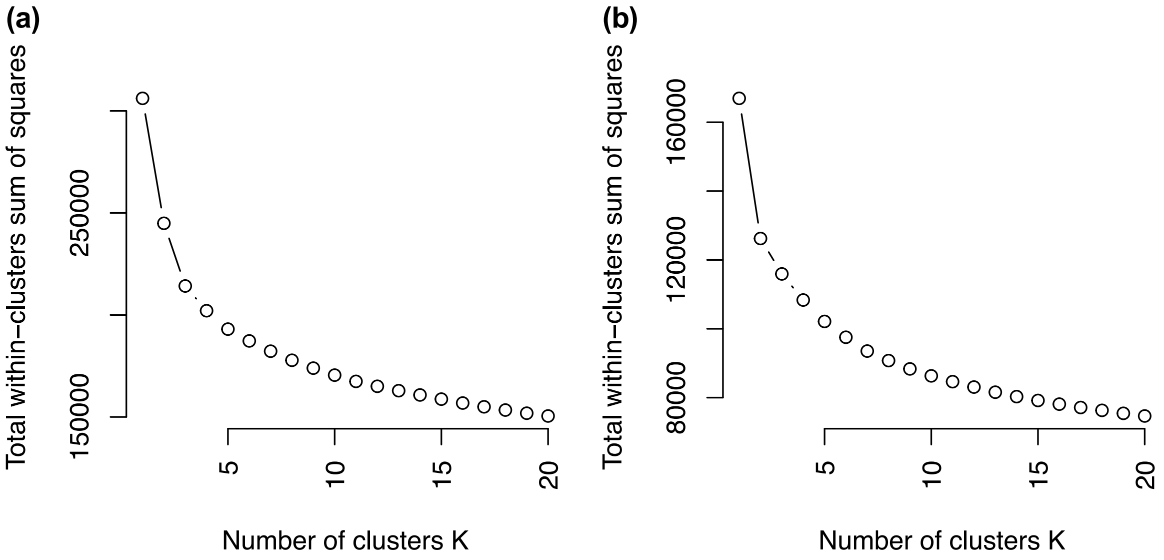


**Figure S2 The elbow method of *k*-means clustering.**
The elbow method of *k*-means clustering was applied to the transcriptome of (a) Red-type and (b) Green-type cultivars.

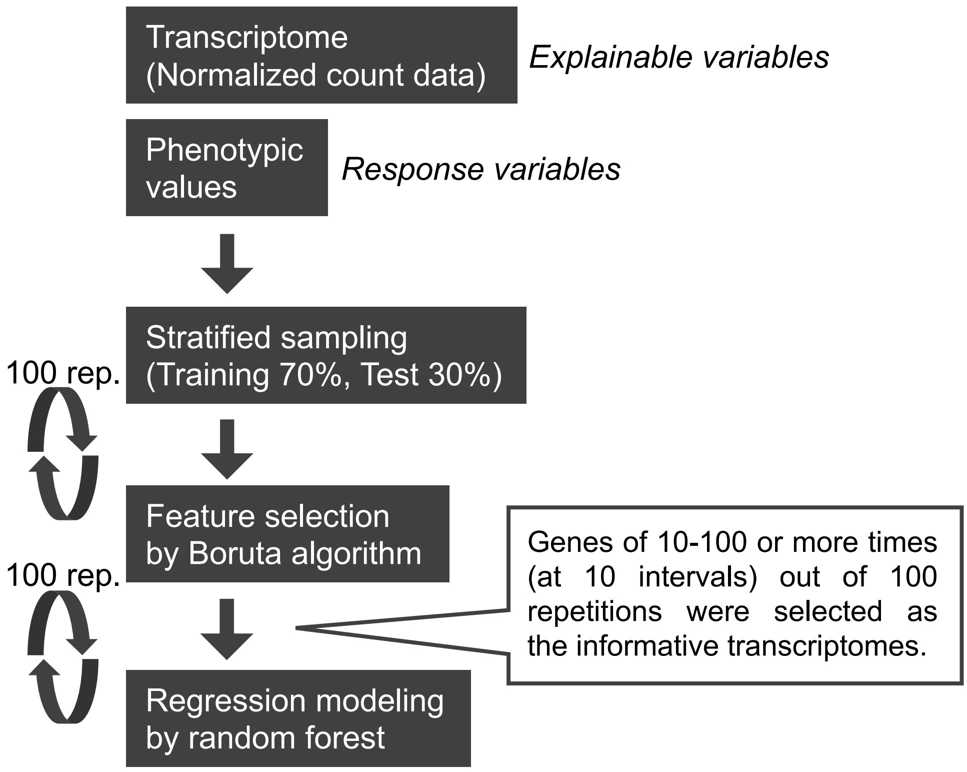


**Figure S3 Flow charts of the regression modeling.**

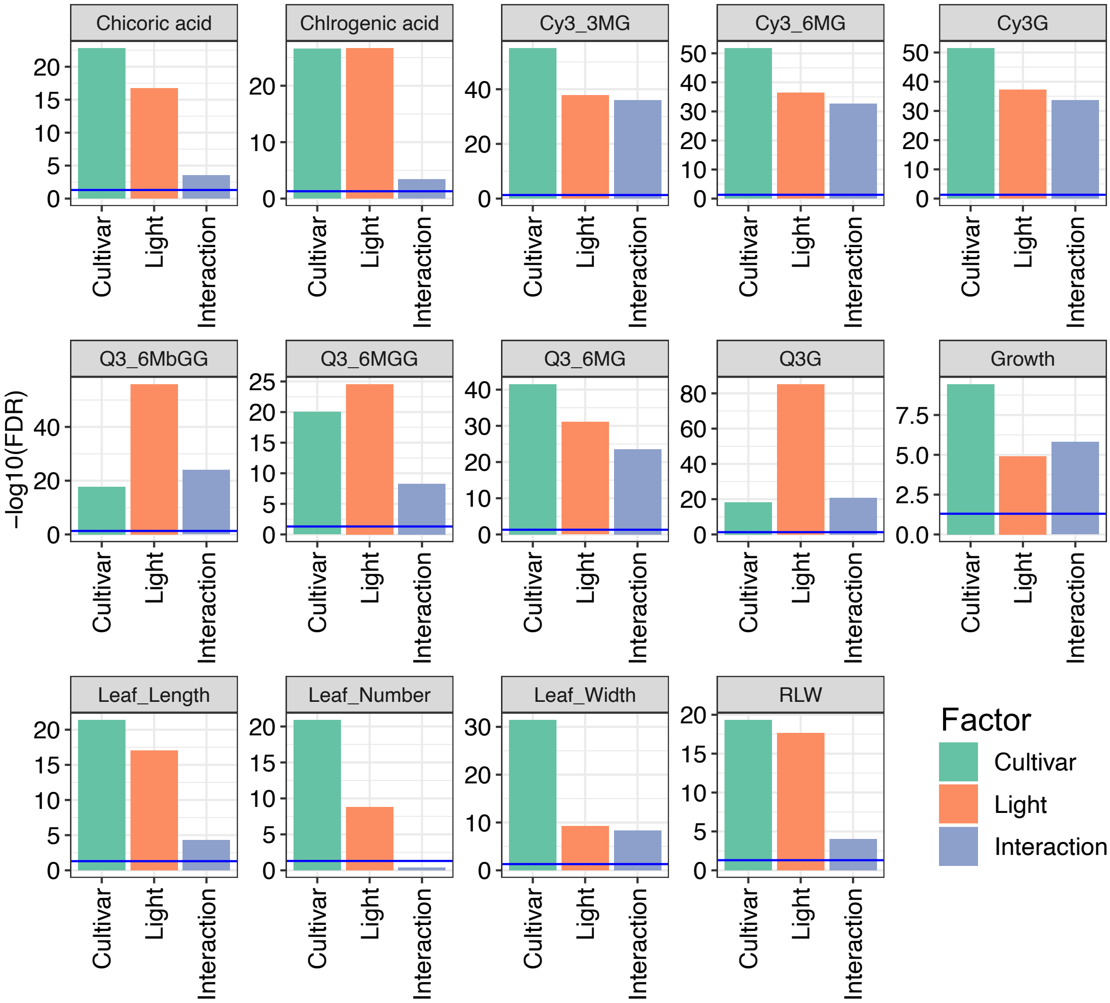


**Figure S4 The statistics of two-way ANOVA for G × E interaction.**
Blue lines indicate the values of $-\log_{10}$ (FDR=0.05).

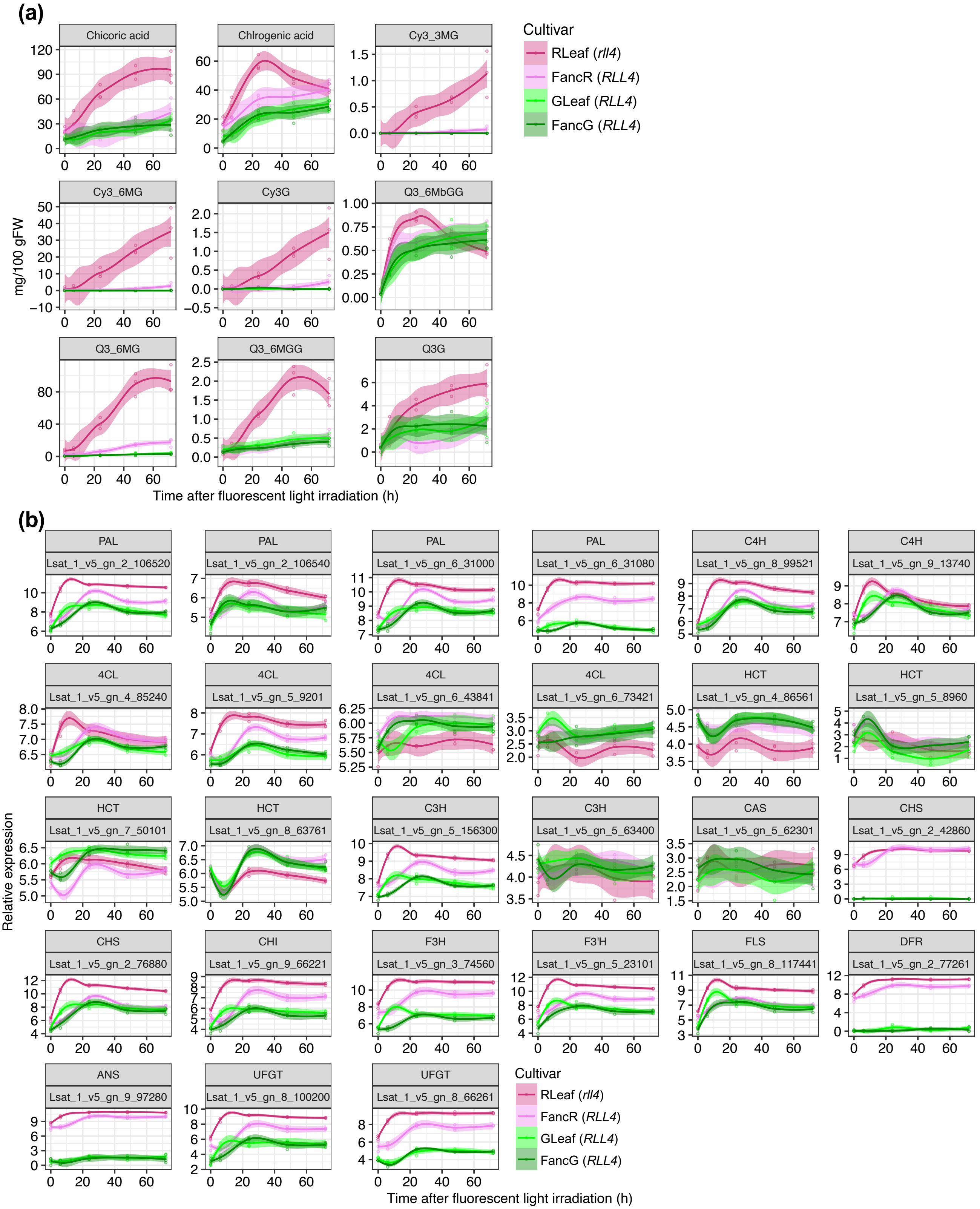


**Figure S5 Time-series response of the phytochemical metabolism in four leaf lettuce cultivars under fluorescent light conditions.**

**(a)** Time-series response of phytochemical contents **(b)** Time-series response of phenylpropanoid and flavonoid pathway gene expressions.

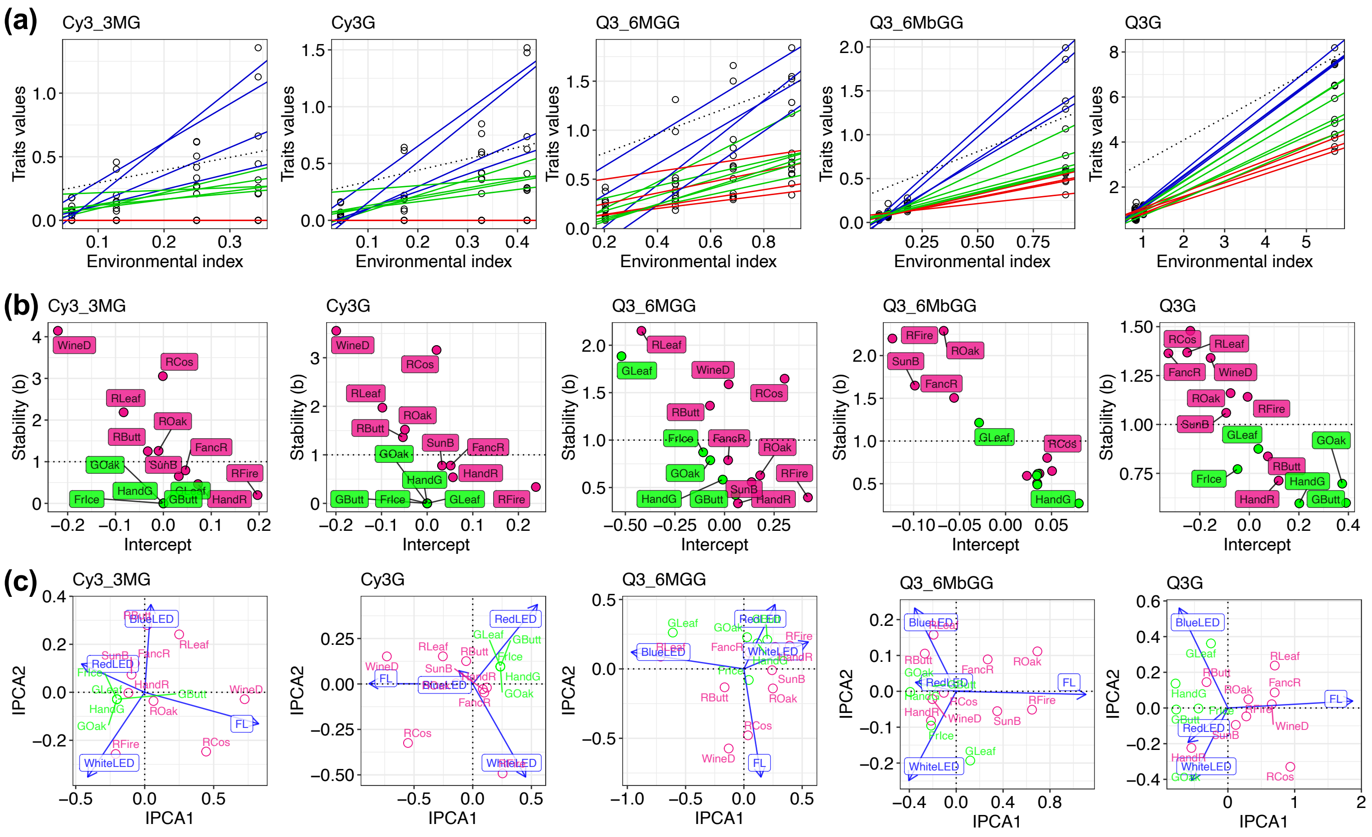


**Figure S6 The analysis of G × E interaction for phytochemical contents.**
**(a, b)** The joint regression analysis (JRA) based on Finlay-Wilkinson model. **(a)** Each plot indicates the observed phenotypic value for each genotype in an artificial light environment, and each line is the regression-fitted value for each genotype. **(b)** The relationship of each stability (slope of JRA), means each slope, and intercept in the regression-line for each genotype. **(c)** Biplots of additive main effects and multiplicative interaction (AMMI) model. The first and second interaction principal component axes (IPCA) were plotted.

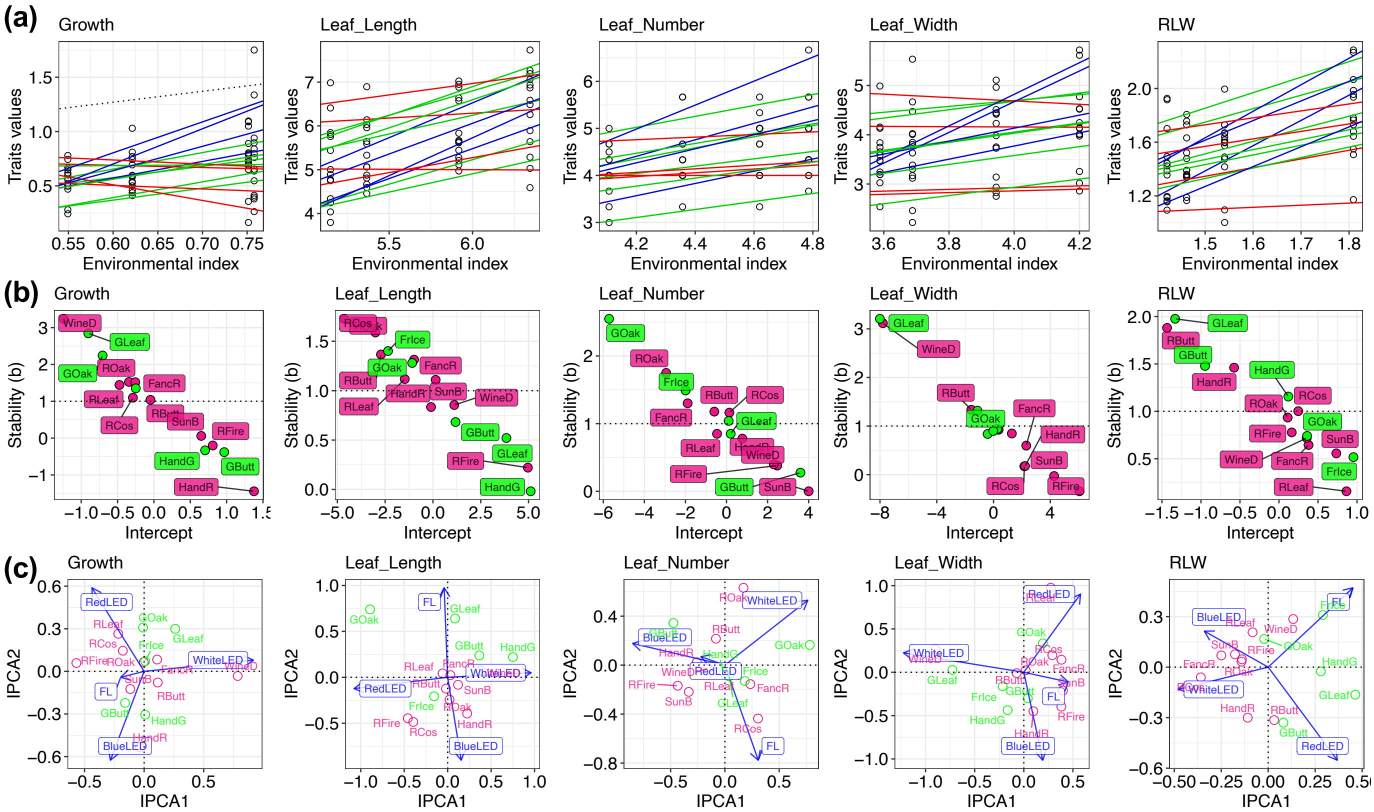

**Figure S7 The analysis of G × E interaction for morphological traits.**
**(a, b)** The joint regression analysis (JRA) based on Finlay-Wilkinson model. **(a)** Each plot indicates the observed phenotypic value for each genotype in an artificial light environment, and each line is the regression-fitted value for each genotype. **(b)** The relationship of each stability (slope of JRA), means each slope, and intercept in the regression-line for each genotype. **(c)** Biplots of additive main effects and multiplicative interaction (AMMI) model. The first and second interaction principal component axes (IPCA) were plotted.

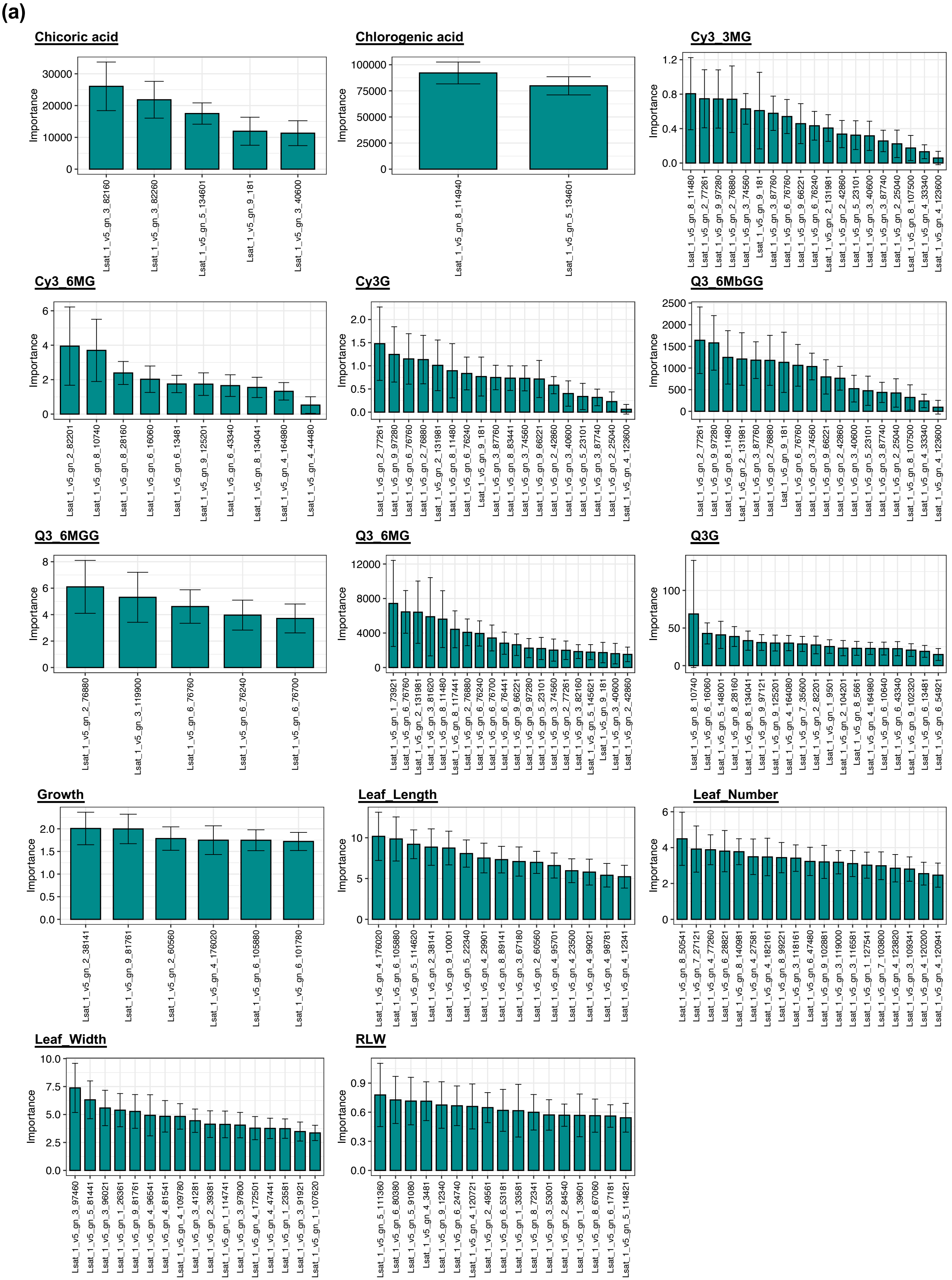


**Figure S8 Interpreting of a transcriptome-based black-box model in all phenotypes.**
**(a)** Variable importance in the regression model based on random forest algorithm for each phenotype. Data and error bars are the mean ± SD (100 repetitions). **(b)** Partial dependence plots (pdp) in the regression model based on random forest algorithm for each phenotype. Blue curves show LOESS smoothing lines of the pdp in 100 repetitions.

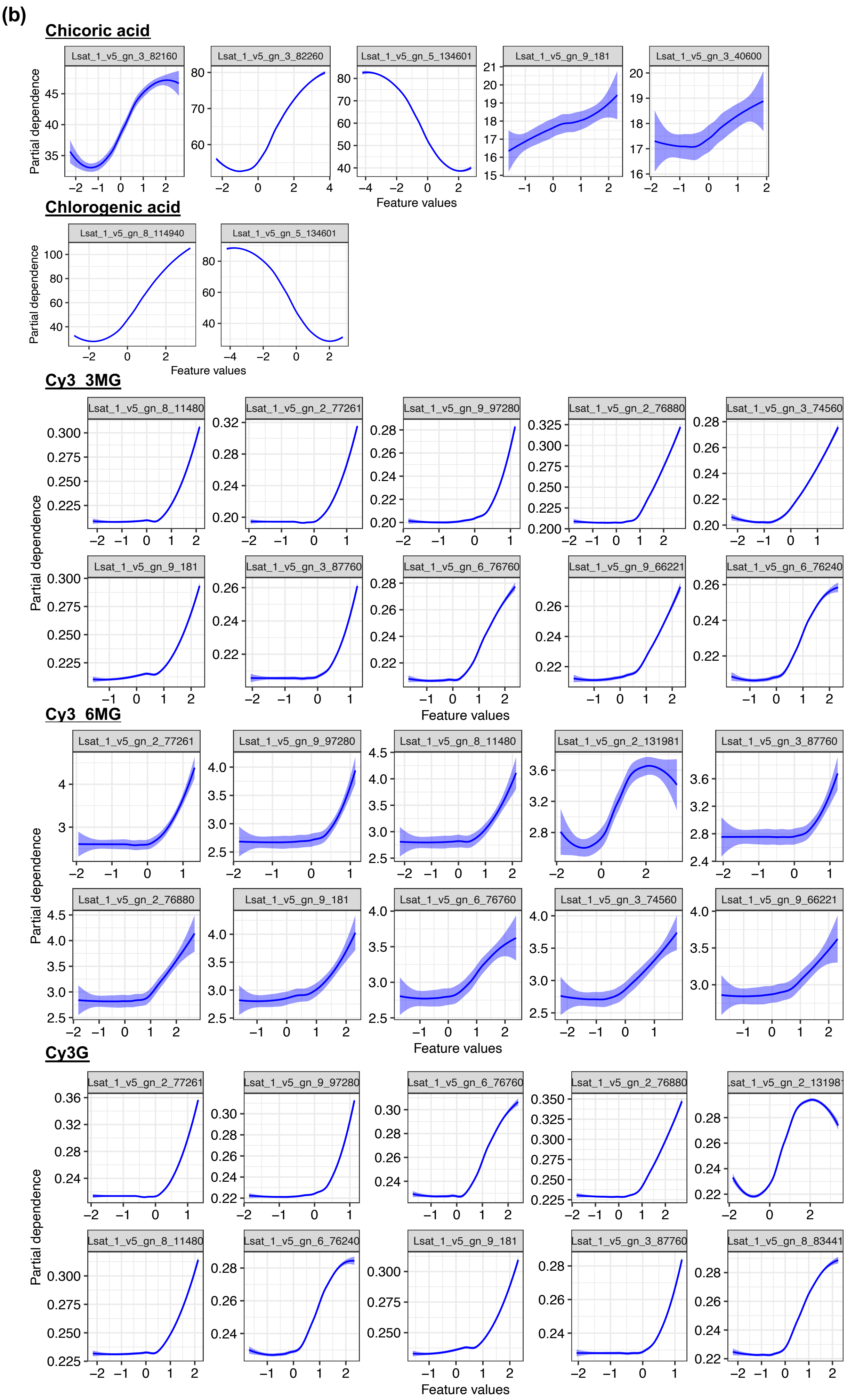


**Figure S8 continued**

(b)

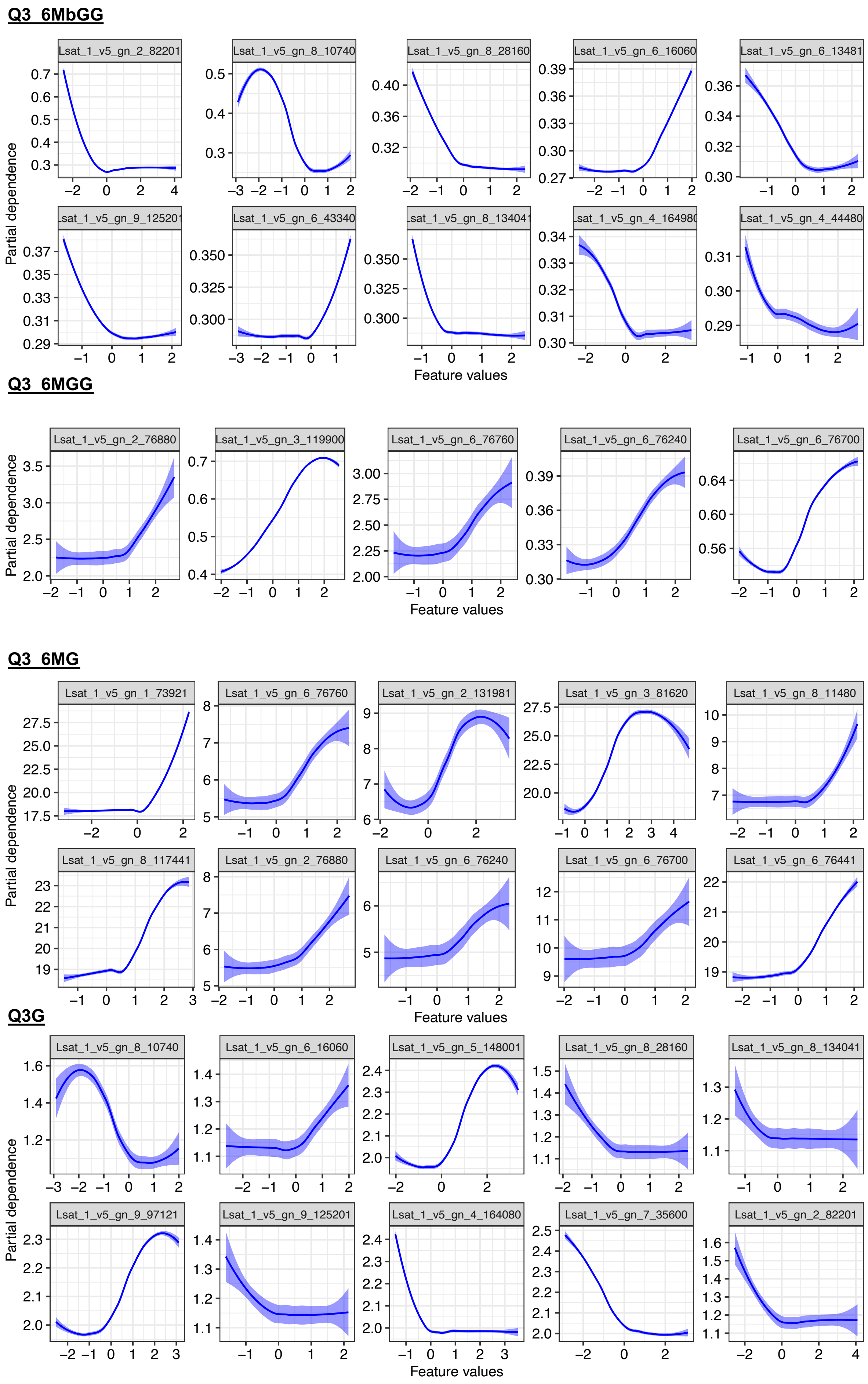


**Figure S8 continued**

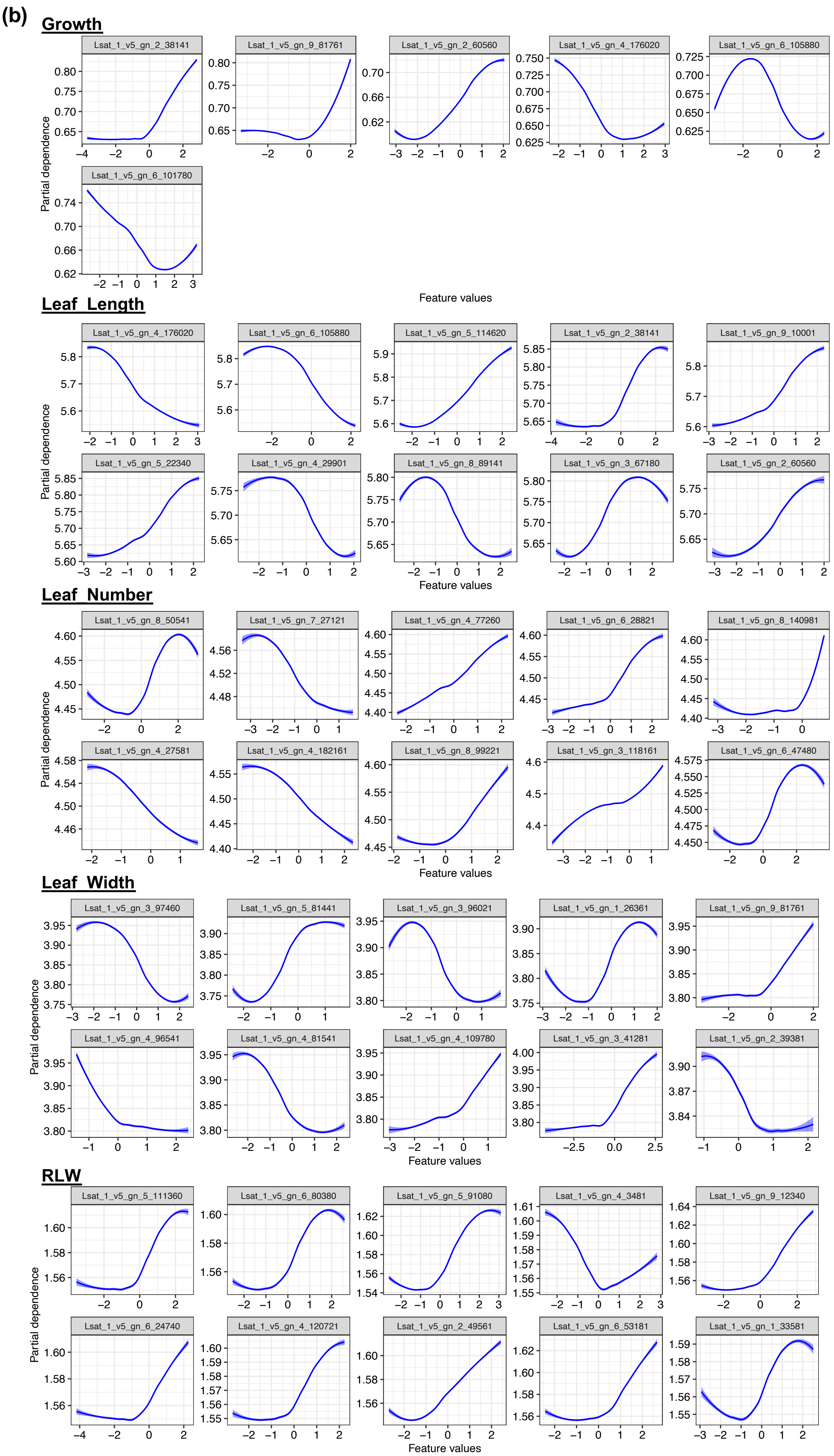


**Figure S8 continued**

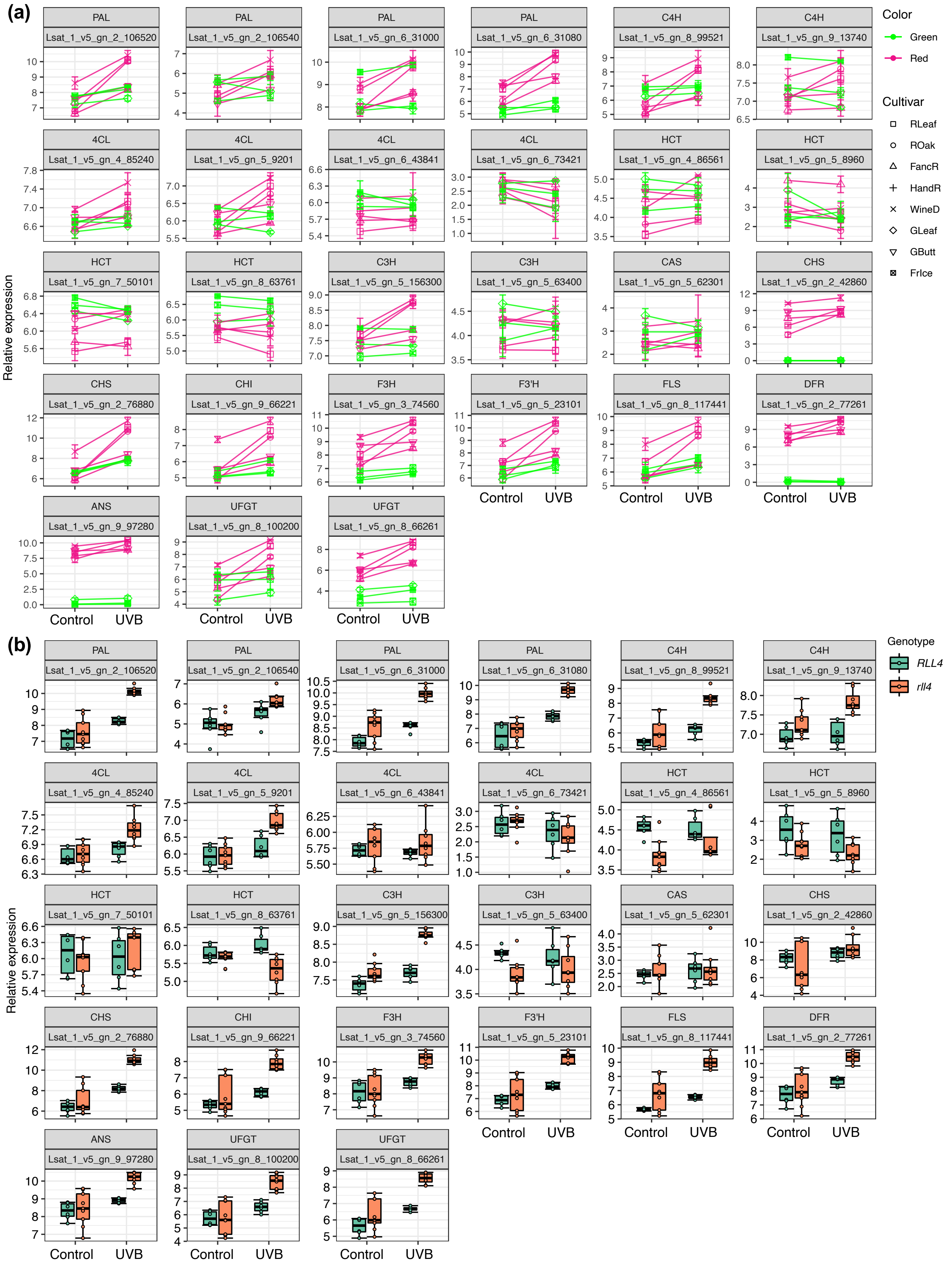


**Figure S9 UV-B response of phenylpropanoid and flavonoid pathway gene expressions in eight lettuce cultivars.**

**(a)** Interaction plots of expression levels. **(b)** Boxplots of expression levels among *RLL4* alleles in red-type cultivars.

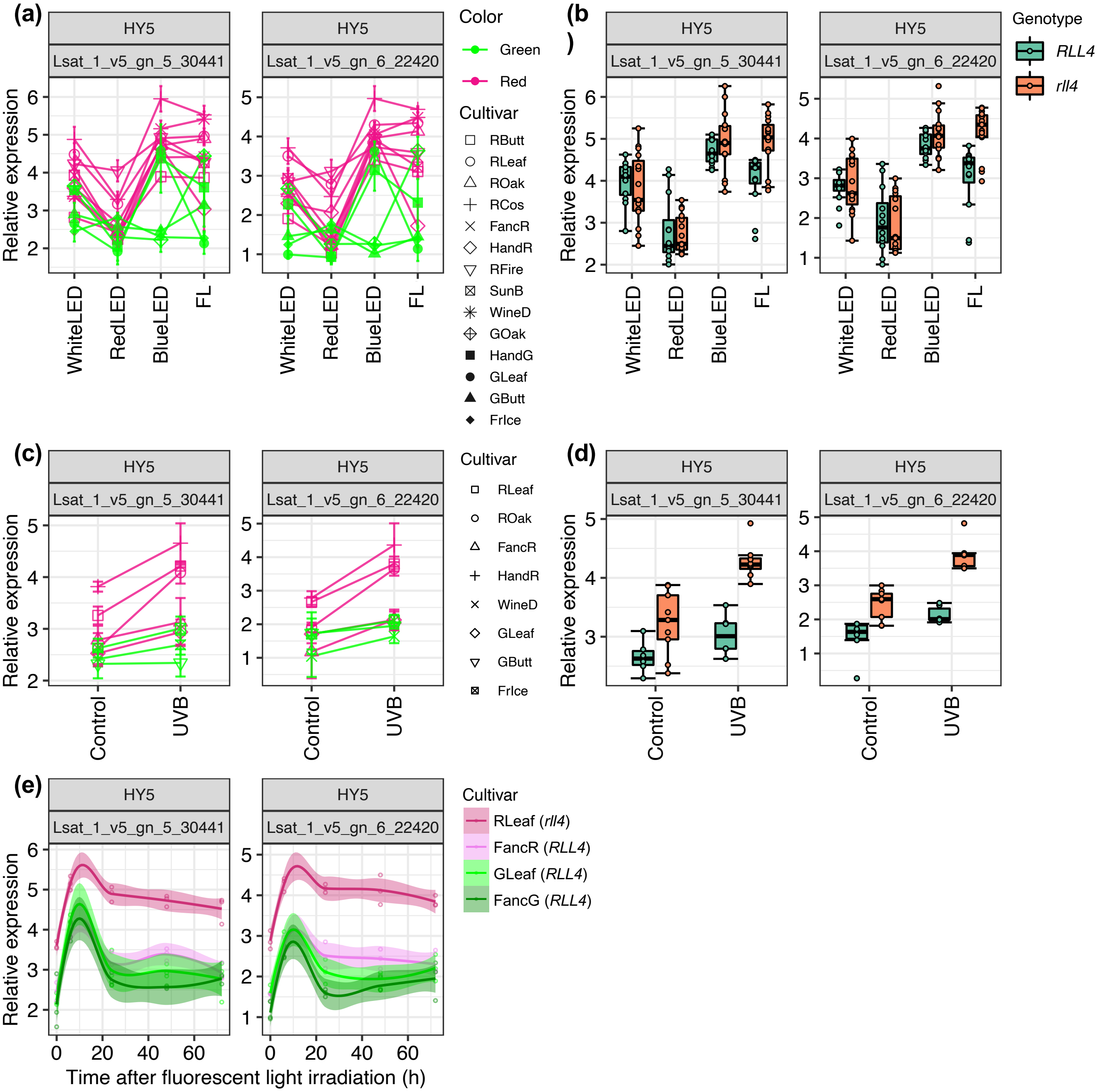


**Figure S10 Expression patterns of lettuce HY5 genes under artificial light conditions.** **(a, c)** Interaction plots of expression levels under four artificial light and UV-B supplementation conditions. **(b, d)** Boxplots of expression levels among *RLL4* alleles in red-type cultivars under four artificial light and UV-B supplementation conditions. **(e)** Time-series response of HY5 gene expressions in four leaf lettuce cultivars under fluorescent light conditions.

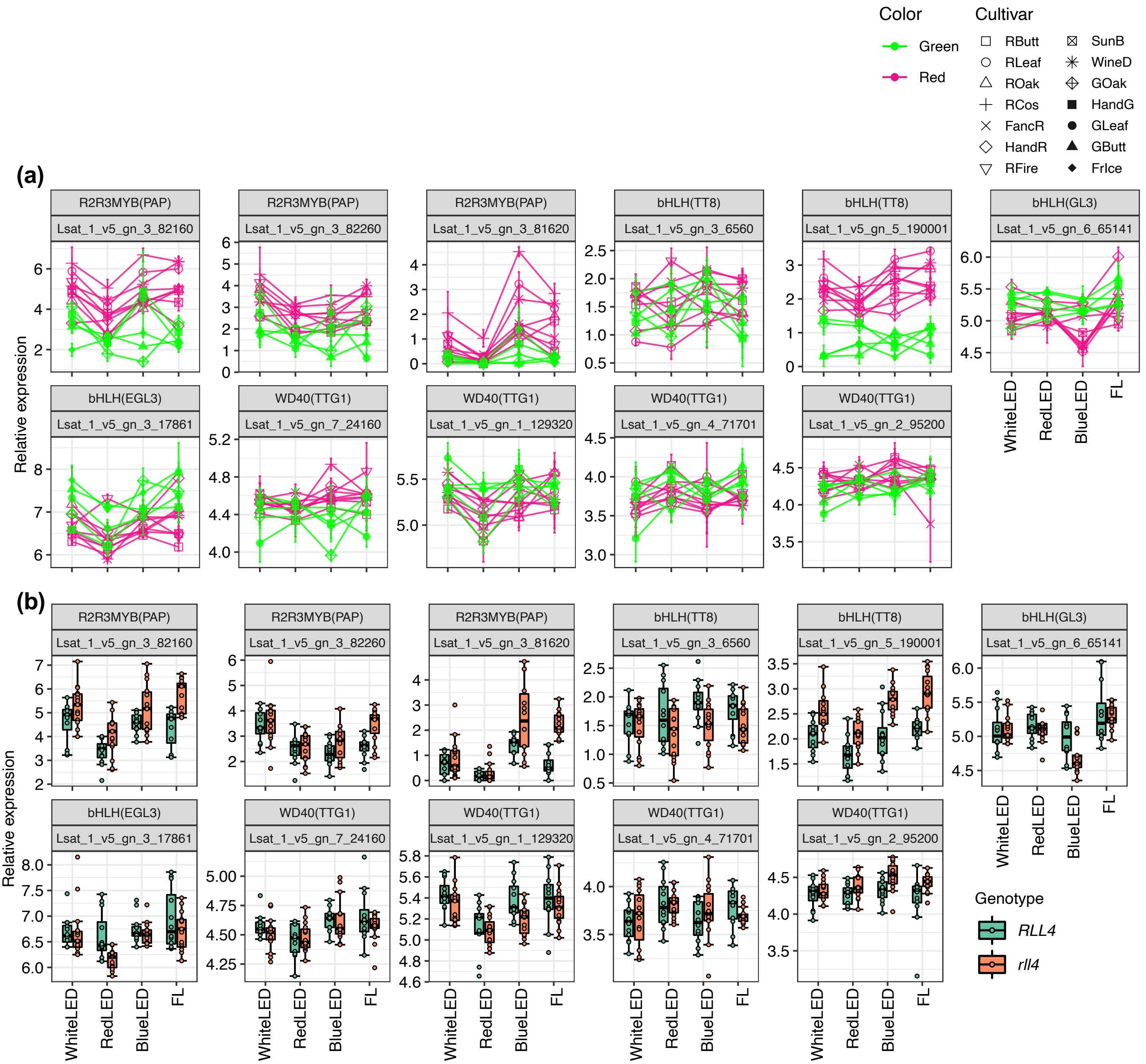


**Figure S11 Expression patterns of lettuce MBW complex genes under artificial light conditions.** **(a, c)** Interaction plots of expression levels under four artificial light and UV-B supplementation conditions. **(b, d)** Boxplots of expression levels among *RLL4* alleles in red-type cultivars under four artificial light and UV-B supplementation conditions. **(e)** Time-series response of gene expressions in four leaf lettuce cultivars under fluorescent light conditions.

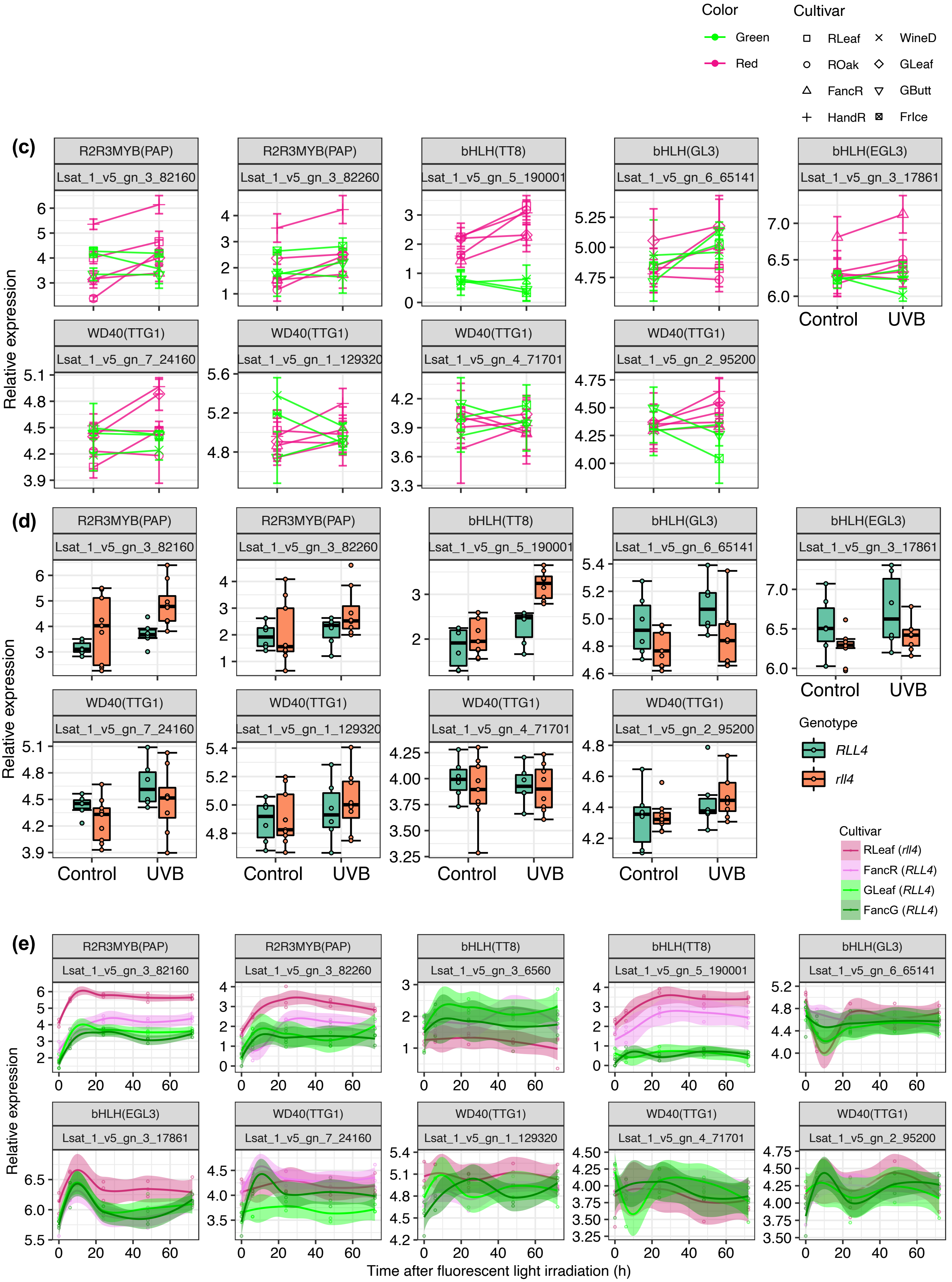


**Figure S11 continued**

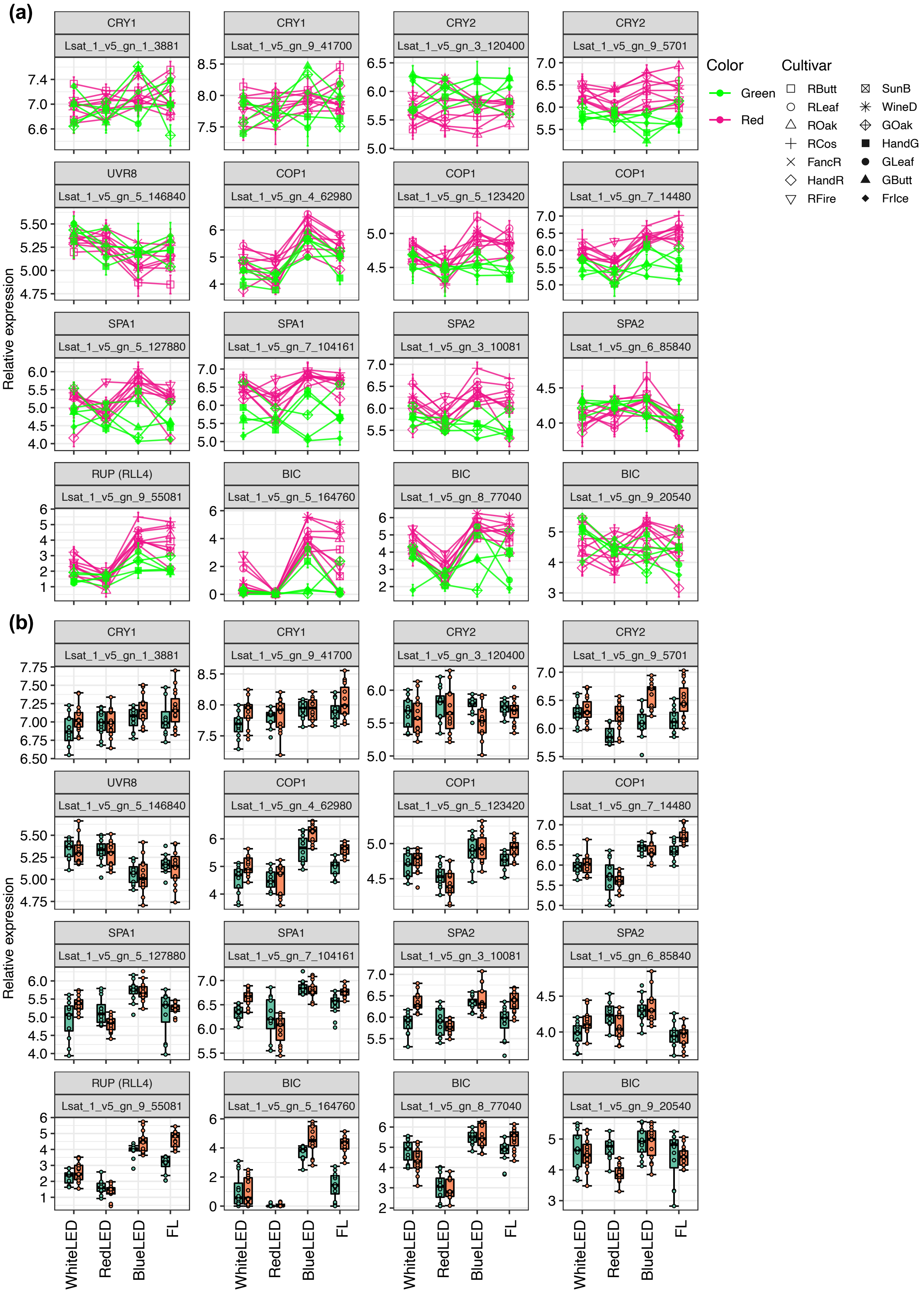


**Figure S12 Expression patterns of lettuce light signal-related genes under artificial light conditions.** **(a, c)** Interaction plots of expression levels under four artificial light and UV-B supplementation conditions. **(b, d)** Boxplots of expression levels among *RLL4* alleles in red-type cultivars under four artificial light and UV-B supplementation conditions. **(e)** Time-series response of gene expressions in four leaf lettuce cultivars under fluorescent light conditions.

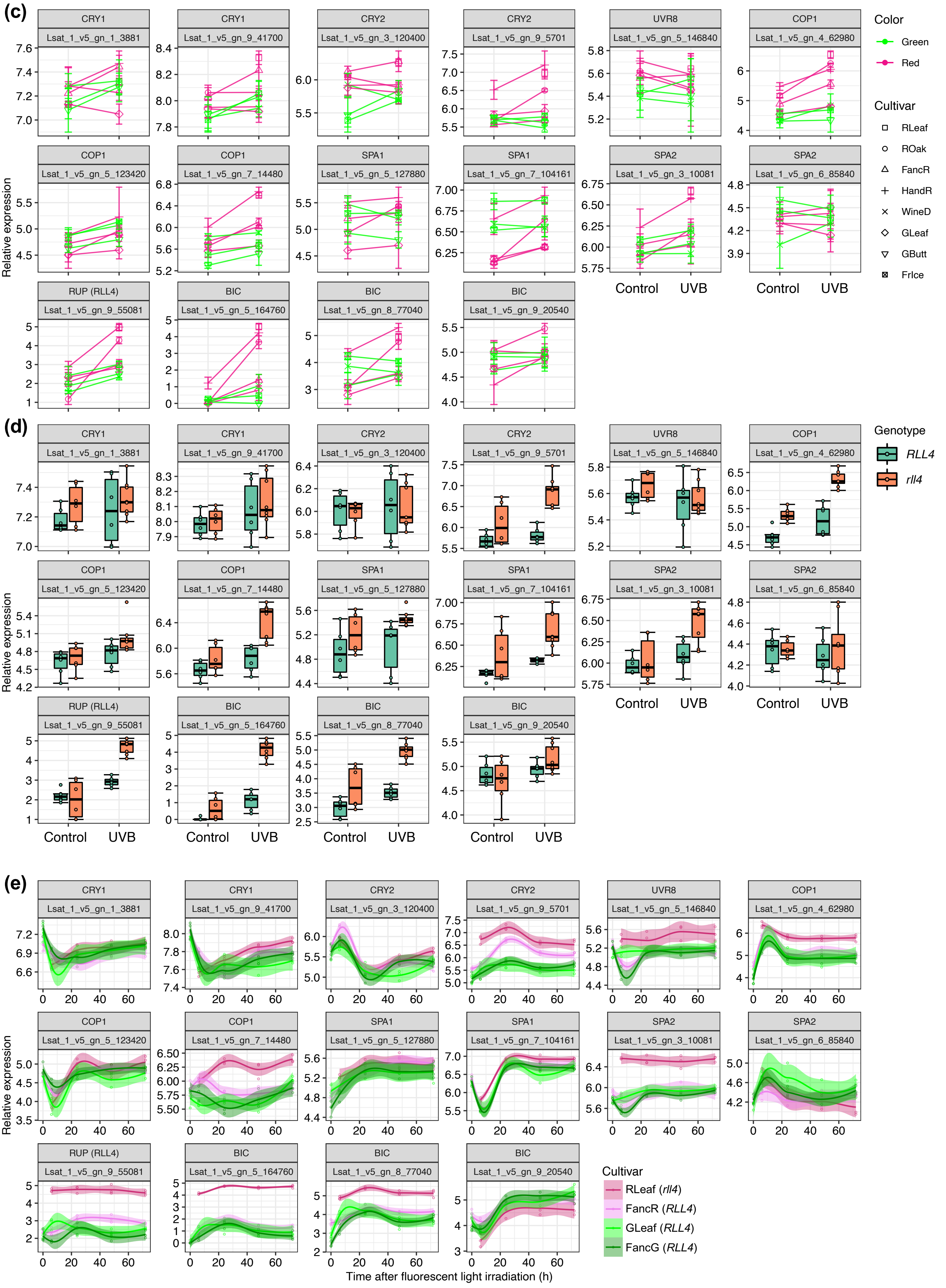


**Figure S12 continued**
